# Meta-analysis of the acoustic adaptation hypothesis reveals no support for the effect of vegetation structure on acoustic signalling across terrestrial vertebrates

**DOI:** 10.1101/2024.02.21.581368

**Authors:** Bárbara Freitas, Pietro B. D’Amelio, Borja Milá, Christophe Thébaud, Tim Janicke

## Abstract

Acoustic communication plays a prominent role in various ecological and evolutionary processes involving social interactions. The properties of acoustic signals are thought to be influenced not only by the interaction between signaller and receiver but also by the acoustic characteristics of the environment through which the signal is transmitted. This conjecture forms the core of the so-called “acoustic adaptation hypothesis” (AAH), which posits that vegetation structure affects frequency and temporal parameters of acoustic signals emitted by a signaller as a function of their acoustic degradation properties. Specifically, animals in densely vegetated ‘closed habitats’ are expected to produce longer acoustic signals with lower repetition rates and lower frequencies (minimum, mean, maximum, and peak) compared to the ones inhabiting less vegetated ‘open habitats’. To date, this hypothesis has received mixed results, with the level of support depending on the taxonomic group and the methodology used. We conducted a systematic literature search of empirical studies testing for an effect of vegetation structure on acoustic signalling and assessed the generality of the AAH using a meta-analytic approach based on 371 effect sizes from 75 studies and 57 taxa encompassing birds, mammals and amphibians. Overall, our results do not provide consistent support for the AAH, neither in within-species comparisons (suggesting no overall phenotypically plastic response of acoustic signalling to vegetation structure) nor in among-species comparisons (suggesting no overall evolutionary response). However, when considering birds only, we found a weak support for the AAH in within-species comparisons, which was mainly driven by studies that measured frequency bandwidth, suggesting that this variable may exhibit a phenotypically plastic response to vegetation structure. For among-species comparisons in birds, we also found support for the AAH, but this effect was not significant after excluding comparative studies that did not account for phylogenetic non-independence. Collectively, our synthesis does not support a universal role of vegetation structure in the evolution of acoustic communication. We highlight the need for more empirical work on currently under-studied taxa such as amphibians, mammals, and insects. Furthermore, we propose a framework for future research on the AAH. We specifically advocate for a more detailed and quantitative characterization of habitats to identify frequencies with the highest detection probability and to determine if frequencies with greater detection distances are preferentially used. Finally, we stress that empirical tests of the AAH should focus on signals which are selected for increased transmission distance.

## I. INTRODUCTION

### (1) Acoustic signalling and the effect of multiple selective forces

Acoustic signalling is an essential facet of animal communication, spanning a wide array of species and ecosystems. These signals play a pivotal role in various contexts, including mate attraction, territory defence, predation avoidance (alerting others of a threat), rival intimidation, individual recognition, and social communication (Catchpole & Slater, 2008; Bradbury & Vehrencamp, 2011). Thus, acoustic signals are subject to multiple selective forces imposed on the signaller and receiver that shape them to finely tuned instruments of communication (Wiley & Richards, 1982; Catchpole, 1987; Catchpole & Slater, 2008). These multiple selective forces may act in concert or in opposition, which potentially complicates our understanding of the evolution of acoustic communication (i.e., the adaptive significance of the specific structure of a given signal).

One of the most studied selective forces shaping acoustic signals is the morphology of the signaller. For instance, the size of sound-producing organs is often positively associated with a signaller’s body size or mass and has been shown to influence signal frequency in anurans and mammals (Fletcher, 2004; Tonini *et al*., 2020). Similarly, in birds body size has been found to be negatively correlated with signal minimum frequency, and positively with frequency bandwidth (Friis *et al*., 2022) or peak frequency (Mikula *et al*., 2021). Thus, when individuals emit sounds, they also convey information about their size such that acoustic cues can serve as honest signals of body condition and reproductive performance, influencing mate choice (McLean, Bishop & Nakagawa, 2012; Welklin *et al*., 2023). Although producing acoustic signals at high amplitude may attract more mates and provide competitive advantages, it incurs energetic costs (e.g., Gil & Gahr, 2002; Ophir, Schrader & Gillooly, 2010), which can make it challenging to produce an optimally designed sound for long-distance transmission. In addition, predators and parasites can use acoustic signals emitted by their prey as cues to locate them (Ryan, Tuttle & Taft, 1981; Garamszegi & Avilés, 2005), potentially driving the development of signals that minimize their detectability, such as high-frequency pure tones in certain alarm calls to reduce locatability (Klump & Shalter, 1984). In well-documented vocal learners such as oscine passerines, hummingbirds, parrots, humans, elephants, seals, and toothed whales, vocalisations are also shaped by the capability of reproducing heard sounds (Tyack, 2020). This ability might counterbalance other predicted trends (mentioned above) that shape acoustic signals. For example, learning different signals from immigrants or learning with errors can introduce variations that offset these trends. Likewise, interactions with other animals can shape acoustic signals in multiple ways, for example by selecting for frequencies that exploit sensory biases (Cummings & Endler, 2018).

The structure of acoustic signals can also be influenced by the acoustic characteristics of the environment into which they are emitted and received (Chappuis, 1971; Slabbekoorn & Smith, 2002; Derryberry *et al*., 2018). While propagating from signaller to receiver, signals are subject to several alterations in temporal and frequency parameters. These alterations include reverberation, i.e., the delay between the faster waves (direct) and slowest waves (indirect) causing distortion of the sound (Morton, 1975; Wiley & Richards, 1978, 1982), and attenuation, i.e., the progressive reduction in amplitude which can be attributed to three main components: (1) geometric attenuation (also known as spreading loss or spherical attenuation), (2) atmospheric attenuation, and (3) habitat attenuation (Embleton, 1996). The latter two components are often collectively referred to as degradation, resulting from scattering, refraction through obstacles, atmospheric absorption, or echoes, with greater effects typically observed for high frequencies (Aylor, 1972; Marten & Marler, 1977). In this context, natural selection is expected to have favoured the evolution of accurate and efficient transmission of acoustic signals for long-distance travel, enabling information transfer from signaller to receiver while accounting for environmental constraints.

### (2) The Acoustic Adaptation Hypothesis

The various selective forces imposed by the environment outlined above form the theoretical background of the acoustic adaptation hypothesis (AAH), mainly grounded in the work of Morton (1975) and Hansen (1979) but coined by Rothstein & Fleischer (1987). The AAH argues that the transmission of acoustic signals is affected by the environment and offers explicit predictions regarding the optimal shape of acoustic signals to ensure accurate transmission in environments characterized by high degradation, such as densely vegetated habitats (Morton, 1975; Blumenrath & Dabelsteen, 2004). Animals inhabiting such hereafter called ‘closed habitats’ (as opposed to less vegetated ‘open habitats’) are expected to have signals of longer duration because this is expected to increase the likelihood of detection and to leverage the amplitude of reverberated sound waves, extending the range of signal propagation (Nemeth *et al*., 2006). To increase transmission distance in closed habitats, acoustic signals should also have lower repetition rates to minimize interference with reverberated waves (Brown & Handford, 2000; Naguib, 2003) and narrower frequency bandwidths to concentrate energy at frequencies less susceptible to attenuation (Wiley & Richards, 1978). Additionally, animals in closed habitats should be prone to emit acoustic signals at lower frequencies including minimum, mean, maximum, and peak frequency (Chappuis, 1971; Morton, 1975; Marten, Quine & Marler, 1977; see Table S1 for the definition and predictions for each acoustic variable).

The AAH is currently considered as a component of the broader Sensory Drive Hypothesis, which predicts that signals, sensory mechanisms, and microhabitat choice evolve together based on the characteristics of the habitat, ambient noise patterns, presence of predators and parasites, and various sensory and physiological factors (Endler, 1992; Cummings & Endler, 2018). Several authors have also extended the AAH to a wider framework of environmental adaptation, covering studies on the effect of environmental noise such as tests of the Lombard effect in the context of urbanization (Job, Kohler & Gill, 2016; Zhao *et al*., 2021). Here, we focus on the AAH in the context of the effect of vegetation structure on acoustic signalling as originally tested by Morton (1975).

Playback transmission experiments using either natural or synthetic sounds have been conducted to assess the physical foundations of the AAH (Wiley & Richards, 1978). In addition, comparisons between species and populations inhabiting environments with different vegetation structure have been performed to test the validity of this hypothesis. While studies testing environmental filtering of sound have provided a robust foundation for the AAH (reviewed in Hardt & Benedict, 2021), its influence on animal acoustic signal production remains unclear. In fact, empirical support for the AAH is highly variable, with many studies supporting the hypothesis (Bertelli & Tubaro, 2002; Nicholls & Goldizen, 2006) whereas many others contest its validity (Hylton & Godard, 2001; Bosch & De la Riva, 2004; Mikula *et al*., 2021). In addition, Hardt & Benedict (2021) uncovered that animal signals are often not better transmitted in a species’ habitat compared to environments they are not adapted to, raising further questions about the generality of the AAH.

In a previous meta-analysis limited to birds, Boncoraglio & Saino (2007) compiled 91 effect sizes from 26 studies of within- (including 12 species) and among-species (spanning 3 orders) comparisons to test for an overall effect of habitat structure (open versus closed) on acoustic variables. The authors found support for the AAH in terms of a lower minimum frequency, maximum frequency, peak frequency, and frequency bandwidth in closed habitats based on a global effect size of Fisher’s z = 0.160, 0.182, 0.346, and 0.230, respectively (Boncoraglio & Saino, 2007). Interval duration was the only variable that did not differ between habitats. Since its publication, this study has served as a key reference in support of the AAH in birds. However, Boncoraglio & Saino (2007) relied on a limited sample size and did not include measurements of mean frequency, total duration, or repetition rate in their analyses, despite clear predictions for these three variables within the framework of the AAH. Furthermore, this study did not explicitly control for potential sources of statistical non-independence beyond shared study identity, such as shared phylogenetic history. The lack of statistical control for non-independence among effect sizes can result in inaccurate conclusions, often leading to spurious statistical significance (Nakagawa *et al*., 2017), and integrating phylogenetic structure into meta-analyses is now considered critical to obtain unbiased results (Lajeunesse, 2009; Chamberlain *et al*., 2012).

Although the AAH was initially formulated based on avian studies, there has been a growing interest in testing this hypothesis in other taxonomic groups. Aiming to examine how widespread the impact of vegetation structure on the evolution of acoustic signals is in vertebrates, Ey & Fischer (2009) qualitatively reviewed 55 studies on birds, anurans and mammals and concluded that the AAH did not seem to be universally valid across taxa. However, the authors claimed that in birds, effects on minimum frequency and frequency bandwidth appeared to align with the predictions of the AAH, whereas relationships with repetition rate, mean, peak, and maximum frequencies did not, which does not fully correspond to the results obtained by Boncoraglio & Saino (2007).

### (3) Assessing the AAH across animals

Despite the large body of research aimed at testing the AAH, it still remains unclear to what extent this hypothesis is valid across acoustic variables in a broad range of animal taxa, and whether contrasting findings from previous studies reflect methodological artifacts and/or actual differences among taxa. Our study aims to fill this gap by conducting a meta-analysis of empirical studies that tested the AAH in the context of airborne acoustic signalling in response to vegetation structure. In this regard, we build on the work by Boncoraglio & Saino (2007) with the aim to provide a robust and comprehensive meta-analysis. More specifically, our main objective is to address two main questions: (1) Does the AAH hold across different animal taxa and acoustic parameters in the context of vegetation structure? and (2) does the AAH support phenotypically plastic and/or evolutionary responses to vegetation structure (inferred from within- and among-species comparisons, respectively)? For these purposes, we compiled 371 effect sizes testing the AAH in animals. We did not have a clear prediction on whether effect sizes differ between within- and among-species comparisons. However, it seems plausible that effect sizes from among-species comparisons might be weaker because acoustic parameters likely evolve not only in relation to the habitat but also due to other factors such as interspecific variation in mate preferences and/or neutral processes such as genetic drift.

## II. METHODS

This meta-analysis was pre-registered at the Open Science Framework (OSF; study details available at: https://doi.org/10.17605/OSF.IO/E7JRS). Differences between the pre-registered study design and the eventually implemented approach are mainly due to methodological improvements and are explained in Text S1.

### (1) Literature search

We conducted a systematic literature search to obtain an unbiased sample of studies testing for a relationship between vegetation structure and acoustic communication in animals following the Preferred Reporting Items for Systematic Reviews and Meta-Analyses approach (PRISMA; Moher *et al*., 2009) in combination with the PRISMA-EcoEvo guidelines (O’Dea *et al*., 2021). We screened the Scopus, the ISI Web of Science Core Collection (Clarivate Analytics), and the Open Access Theses and Dissertations (OATD; https://oatd.org/) databases on 16th December 2021 with the search terms defined as (((“*song” OR “vocal*”) OR ((“acoustic$” OR “bioacoustic$”) AND “call$”)) AND (“habitat$” OR “vegetation”)) OR “acoustic adaptation”. These searches yielded 3300, 3233, and 372 records, respectively. We also considered studies included in previous reviews on related questions (Ey & Fischer, 2009; Hardt & Benedict, 2021) and a previous meta-analysis on birds (Boncoraglio & Saino, 2007), resulting in the addition of 58 studies to be screened. Furthermore, we posted a request for unpublished data on the ‘evoldir’ mailing list (http://life.mcmaster.ca/evoldir.html), the ResearchGate platform (https://www.researchgate.net/), and Twitter, and directly asked colleagues who work on these topics. This resulted in one additional record (Melo *et al*., in prep). Finally, we added one unpublished work on the La Palma population of the Canary Islands Chaffinch (*Fringilla canariensis palmae*; Freitas *et al*., in prep.).

We excluded 45 studies from the OATD search because full texts were not accessible. We also excluded duplicates by following an R workflow adapted from Lagisz (2022) and by manually checking the results of the Zotero deduplication algorithm (Corporation for Digital Scholarship & Roy Rosenzweig Center for History and New Media, 2022). Then, we assigned screening efforts among BF, BM, and TJ, who collectively screened the titles and abstracts of 4558 studies. To ensure consistency, 212 of these studies were screened in parallel by two authors. We selected potentially relevant primary studies using the *metagear* package (Lajeunesse, 2016) based on the following selection criteria:

1. The focus of the empirical study was on non-human animals, for which acoustic signals are transmitted through the air. We did not consider studies focusing on domesticated species as artificial selection can obscure the selection mechanisms that shape communication signals in the wild. We also did not consider studies focusing on echolocation because it is used for orientation in space and/or to locate items rather than for communication and this could confound our results. For syntheses on the effect of vegetation structure on echolocation, see Ey & Fischer (2009). We also excluded studies on aquatic habitats due to the different acoustic properties and difficulties in defining vegetation structure based on underwater vegetation.
2. At least one temporal or frequency parameter for which the AAH predicts an effect was measured for adult individuals of the same species (within-species comparisons) or multiple species (among-species comparisons) across different vegetation types. Studies testing for differences among subspecies were considered as among-species comparisons. Studies that did not replicate habitat type within species or subspecies were excluded (as a conservative approach to mitigate the possibility of testing subspecies differences rather than differences due to habitat). For example, a study was excluded if testing for acoustic signal differences between a subspecies inhabiting one location of habitat type A and another subspecies in another location of habitat type B.
3. Habitat was described based on a continuous variable or on a categorical classification of vegetation structure (from open to closed habitats). If the categorical classification of the habitat was done in more than the two categories, we pooled different habitat types into two categories: ‘open’ and ‘closed’ (e.g., grassland, open field, and meadow grouped in ‘open habitat’). We did not include studies examining differences between urban and rural habitats.
4. The study was written in English, French, German, Portuguese, or Spanish according to the languages that the authors were able to understand.
5. The study reported sufficient information to compute effect sizes (e.g., sample size, test-statistic, means with estimates of uncertainty).

During the title and abstract screening stage, we excluded 4237 studies thatdid not fit at least one of the requirements listed above. Afterwards, we carried out a full-text screening of the remaining 321 studies. During this stage we excluded 246 studies because they: (i) did not test the influence of vegetation structure on acoustic variables (N = 116), (ii) used habitat types that could not be classified as ‘open’ or ‘closed’ (N = 17), (iii) did not replicate habitat type (N = 5), (iv) focused on echolocation (N = 23), (v) focused on playback/transmission experiments (N = 15), (vi) focused on the effect on urbanization (N = 4), (vii) used acoustic variables for which the AAH does not generate predictions (N = 3), (viii) did not report sufficient data (N = 27; i.e., multivariate statistics not providing raw data, missing key data required to compute an effect size even after contacting the authors), (ix) represented reviews or non-empirical studies (N = 14), (x) were inaccessible (N = 12), (xi) were written in other languages (N = 4), or (xii) represented duplicates (N = 6; of which 5 represented theses that are already covered by other primary studies). The final dataset included 75 primary studies (see Fig. 1 for the PRISMA diagram and Table S2 for a reference list of all primary studies).

**Fig. 1.**
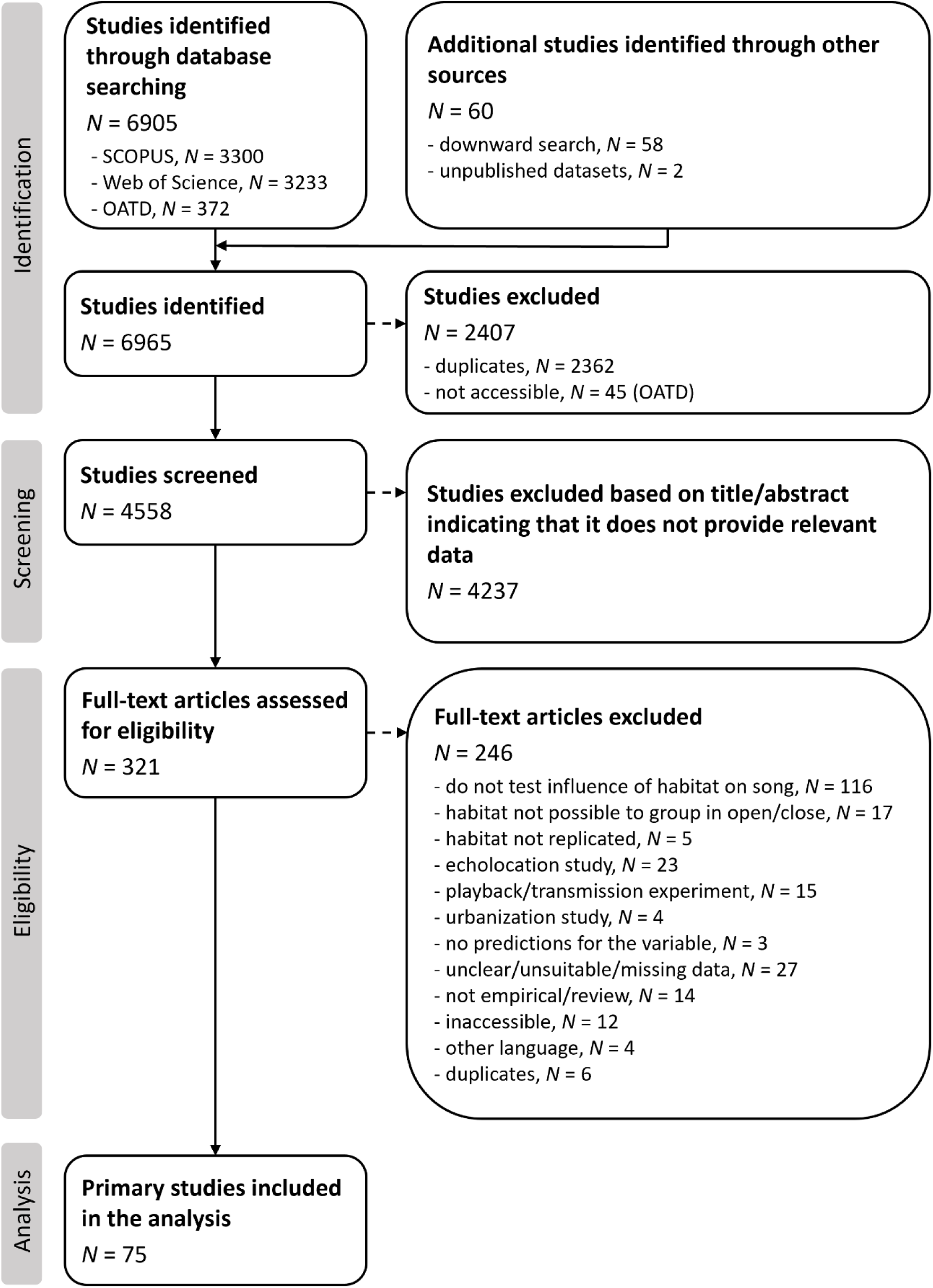
Preferred Reporting Items for Systematic Reviews and Meta-Analyses (PRISMA) diagram of the protocol for screening, including and excluding studies for this meta-analysis and number of studies identified at each stage. OATD, Open Access Theses and Dissertations.

## (2) Data collection

BF and TJ extracted means, metrics of uncertainty (standard deviation or standard error) and sample sizes, correlation coefficients and sample sizes, or the statistical test value and degrees of freedom reported for the relationship among acoustic variables and vegetation structure from the text, tables, figures or supplementary materials. We used the R package *juicr* version 0.1 (Lajeunesse, 2021) to extract summary statistics from figures. From five studies that used multivariate statistics (such as principal components analysis), we reanalysed the raw data if published together with the article or provided by the authors (*N* = 2), and corrected also for the phylogeny in cases in which phylogenetic data was provided (*N* = 1). Extractions from 27 studies were cross-checked by both authors to ensure accuracy.

All comparisons included in our meta-analysis followed a descriptive approach, meaning that none of the primary studies manipulated vegetation structure experimentally to test for a causal effect on acoustic variables. Thus, we used the Pearson correlation coefficient (*r*) as an effect size to synthesize the findings of primary studies. We converted test statistics provided in the primary studies into *r* and estimated its sampling variance (*Vr*) using formulas and guidelines reported in Lajeunesse *et al*. (2013) and using the R package *esc* version 0.5.1 (Lüdecke, 2019). To improve transparency and allow for reproducibility of data extraction, we report the method used to compute effect sizes (Table S2) following a framework provided in Ivimey-Cook *et al*. (2023). For each acoustic variable, correlations that aligned with the predictions of the AAH (Table S1) were assigned a positive sign. For studies using a categorical classification of the vegetation structure (see section ‘Moderator variables’) of three groups (open, closed, and mixed; *N* = 3) and where effect sizes were computed from group means, we only used the values from the two extreme categories: open and closed. All effect sizes were assigned to a type of study: within- or among-species comparisons, to allow testing for a phenotypically plastic or an evolutionary response of acoustic signalling to vegetation structure, respectively. Noting that most of the studies on within-species comparisons focused on nearby populations, we interpreted the support for the AAH in within-species comparisons as suggestive of a phenotypically plastic response rather than adaptive response since we assumed no genetic divergence between the different populations tested within any given species.

### (3) Moderator variables

Besides the global test for the support of the AAH across animals, we aimed at explaining heterogeneity in effect sizes among studies from a biological and methodological perspective. Specifically, we explored if support for the AAH differed among animal taxa and acoustic variables. If an acoustic variable was measured for parts and for the full signal duration, we extracted both effect sizes separately but assigned them as the same acoustic variable (e.g., song duration and note duration were both considered as duration).

In addition, we tested for four methodological moderators: habitat classification, habitat contrast, body size correction, and phylogeny correction. Firstly, we explored effects of the habitat classification (continuous versus categorical) used in primary studies, i.e., if the studies used a continuous variable or a categorical classification to describe the vegetation structure of the habitat. When authors of primary studies ran analyses with both types of classification (*N* = 2), we retrieved only the estimates obtained for the continuous variable, as we consider it less subjective than a categorical classification. Secondly, we scored the magnitude of contrast between closed and open habitats (for studies using a categorical classification) that were used in the study (habitat contrast), attributing a ‘0’ when there was a moderate difference in vegetation structure (e.g., temperate broadleaf forest versus tropical rainforest), or a ‘1’ when there was a substantial difference in vegetation structure (e.g., grassland versus forest). Importantly, primary studies investigated various habitat types and used different methods for classifying them. This diversity in methods to classify vegetation structure could have contributed to differences in study outcomes (suggested by Hardt & Benedict, 2021). By considering these two moderators, we tried to account for this potential source of heterogeneity among studies. Thirdly, we recorded whether primary studies accounted for body size differences within and/or among species, since this trait is known to be allometrically correlated with the morphology of the sound apparatus, and therefore to affect the production of sound (Pearse *et al*., 2018). Finally, we recorded whether studies reporting among-species comparisons accounted for phylogenetic non-independence.

### (4) Phylogeny

In order to account for phylogenetic non-independence and to quantify the phylogenetic signal (also termed phylogenetic heritability; de Villemereuil & Nakagawa, 2014), we reconstructed a phylogeny of all sampled taxa from studies of within- (Fig. S1) and among-species comparisons (Fig. S2). We used the R package *rotl* (Michonneau, Brown & Winter, 2016) to retrieve the phylogenetic relationships from the Open Tree of Life (opentreeoflife.org/; Hinchliff *et al*., 2015) and *ape* (Paradis, Claude & Strimmer, 2004) to obtain the branch lengths of the trees by applying Grafen’s method (Grafen, 1989).

In analyses of among-species comparisons, the phylogenetic position of some taxonomic groups could not be assessed unequivocally so that we assigned the group in question to a higher taxonomic level. Specifically, we considered Parulinae as Parulidae and Cardinalidae as Passeriformes. This decision accounts for the fact that a number of genera traditionally placed within Cardinalidae are embedded within other families, and conversely, some birds traditionally placed in other families belong with the cardinals (Winkler, Billerman & Lovette, 2020). Note that some primary studies tested the AAH across higher taxonomic ranks (such as Anura, Aves and Passeriformes), while others tested it across a lower taxonomic rank, which sometimes was nested within the higher ones (artificial pairs, e.g., Anura and Hylidae, Aves and *Corvus*). In such cases, we retrieved pairwise estimates of divergence times for the taxonomic ranks that were not nested from the TimeTree database (http://timetree.org/; Kumar *et al*., 2017). Then, for the artificial pairs, we attributed a mean estimate between the known divergence time estimates of the nested pairs and the ones more distantly related. For example, for the pair Anura - Hylidae, we computed the divergence time as the mean of the difference between the estimate of Hylidae - *Rhinella* (a genus within Anura) and the estimate of Anura - Felidae, since Felidae is the closest outgroup of Anura in our phylogeny. Then, we computed branch lengths using the methods already specified for the within-species comparison.

### (5) Data analyses

All analyses were performed in R 4.2.2 (R Core Team, 2022) through RStudio 1.4.1106 (RStudio Team, 2021), using the packages *metafor* (Viechtbauer, 2010) for the meta-regression models, *brms* (Bürkner, 2017) for models with Bayesian approach, *orchaRd* (Nakagawa *et al*., 2023a) for calculation of heterogeneity of Restricted Maximum Likelihood (REML) models, *outliers* (Komsta, 2005) for outlier analyses, *dplyr* (Wickham *et al*., 2022) for data manipulation, *ggplot2* (Wickham, 2016) for data visualization, and *ggtree* (Yu *et al*., 2017) to produce our final phylogenies as shown in Figs S1 and S2. Silhouettes in figures were extracted from ‘phylopic’ (https://phylopic.org) using *rphylopic* (Gearty, Jones & Chamberlain, 2023). All data, code, and package versions used are available at https://osf.io/b8km9.

#### (a) Meta-analyses

To provide a global test of the AAH across animals and to assess the impact of moderator variables, we ran both frequentist General Linear-Mixed Effects Models (GLMMs) using REML method and the corresponding Bayesian models. We consider both approaches as complementary, with the Bayesian method having the advantage of estimating the precision of the obtained measures of heterogeneity (Harrer *et al*., 2021). However, for brevity, we report the model results of the Bayesian approach only in the Supporting Information. Bayesian models were run with 4 chains, each with 4000 iterations, a warmup of 2000 iterations, and were checked for convergence. We ran all models for datasets of within- and among-species comparisons separately. First, we obtained a global effect size (*r*) from models in which the effect size obtained from primary studies is defined as the response variable while weighted by the inverse of its sampling variance and with study identifier, effect size identifier, taxon name and the phylogeny (transformed into a correlation matrix) as random effects to account for nonindependence (including no moderators as fixed predictor terms). This was done because the final dataset included multiple effect sizes from species with shared phylogenetic history, from the same species and/or from the same study. Thereby, global effect sizes are estimated while giving more weight to more precise estimates (i.e., those with smaller *Vr*). To correct for non-independence of sampling variances of effect sizes (*Vr*) in REML models, such as shared control statistics, we constructed a variance-covariance matrix quantifying the non-zero covariance between sampling errors within the same primary studies, following the methods by Nakagawa et al. (2023b). We imputed this matrix by assuming that sampling errors within the same study are correlated with a constant ρ = 0.5. Additionally, we checked the robustness of our study results against different values of ρ (0.3, 0.7, and 0.9). Second, each null-model (i.e., including no fixed predictor term) was fitted separately for each taxonomic class (i.e., Aves, Mammalia, Amphibia). Third, we assessed if biological and methodological moderators impacted effect sizes. Specifically, we ran models with the same random factors as above and defined acoustic variable (minimum frequency, mean frequency, maximum frequency, frequency bandwidth, peak frequency, duration, interval duration, and repetition rate), habitat classification (categorical versus continuous), habitat contrast (0 or 1), body size correction (yes or no), or phylogenetic correction (yes or no) as fixed factors. The impact of all moderators was tested separately within birds, which represents the taxonomic class with the highest number of effect sizes (see Results).

We assessed heterogeneity *I*^2^ by estimating the proportion of variance among effect sizes that can be attributed to the different levels of random effects (Higgins & Thompson, 2002). Specifically, we partitioned total heterogeneity (*I*^2^ Total) into variance arising from study identity (*I*^2^ Study), observation (*I*^2^ Observation), taxon (*I*^2^ Taxon), and phylogenetic relationship (*I*^2^ Phylogeny; Nakagawa & Santos, 2012), which is also known as phylogenetic heritability (*H*^2^), and equivalent to Pagel’s λ (de Villemereuil & Nakagawa, 2014). For REML models with predictor variables, we calculated the pseudo-R² based on the proportional change in the sum of the variance components, and its 95% confidence interval based on 1000 bootstrap replicates (Viechtbauer, 2023). For models run with the Bayesian approach, we computed the proportion of variance explained by the entire model (conditional R^2^) and by fixed factors only (marginal R^2^; Nakagawa & Schielzeth, 2013).

#### (b) Sensitivity analyses and publication bias

We explored the robustness of our results by running several sensitivity analyses. Firstly, we identified potential outliers in effect sizes using Grubbs’ test (Grubbs, 1950). Secondly, we verified if any individual studies influenced the results of our analysis disproportionally by conducting a leave-one-out analysis as described by Nakagawa *et al*. (2023b). This method involves repeatedly fitting the model to the dataset with one study removed at a time. Thirdly, we checked whether conclusions remained when using Fisher’s *z* as effect size. In comparison to Pearson’s correlation coefficient *r*, the sampling variance of Fisher’s *z* statistics depends only on the sample size but not on the effect size itself. Thus, we transformed our effect size *r* into *z* and computed its sampling variance using guidelines by Borenstein *et al*. (2009).

In order to assess publication bias, we ran a multilevel meta-regression as suggested by Nakagawa *et al*. (2022). We ran a GLMM with Fisher’s *z* defined as response variable, its standard error, moderators, and year as fixed effects and study identifier, observation identifier, taxon name and the phylogenetic correlation matrix as random factors, which is advised when heterogeneity is high (see Results; Nakagawa *et al*., 2022). This aimed at testing (i) whether the effect size depends on its standard error, which, similarly to the Egger’s regression (Egger *et al*., 1997), may indicate that small studies get published only (or more often) if effect sizes are large enough to provide statistically significant support for the tested hypothesis (small-study effect), and (ii) for time-lag bias (Boutron *et al*., 2019; also known as the bandwagon effect), which suggests that supportive results get published easier in a newly emerging field but as more data are collected and scepticism about the theoretical foundations may arise, non-intuitive results increase (e.g., Jennions & Møller, 2002). As complementary, we used Kendall’s rank correlation test (Begg & Mazumdar, 1994) to quantify funnel plot asymmetry. Studies with low precision and/or high sampling error that do not support the tested hypothesis might be more prone to go unpublished. Consequently, the lack of studies meeting these criteria is expected to drive funnel plot asymmetry (Sterne *et al*., 2011).

## III. RESULTS

### (1) Dataset summary

In total, we obtained 371 effect sizes from 75 studies and 57 taxa encompassing vertebrate taxa only, i.e., amphibians, birds and mammals (Fig. 2 and Table S3). Of these, 219 effect sizes were from studies reporting within-species comparisons (37 species, including 32 birds, 3 mammals, and 2 amphibians, with 189, 16, and 14 effect sizes, respectively) and 152 were from studies of among-species comparisons (20 taxa, including 14 birds, 3 mammals, and 3 amphibians, with 129, 11, and 12 effect sizes, respectively; a more detailed overview of the sample sizes with respect to the tested moderators is provided in Table S3).

**Fig. 2.**
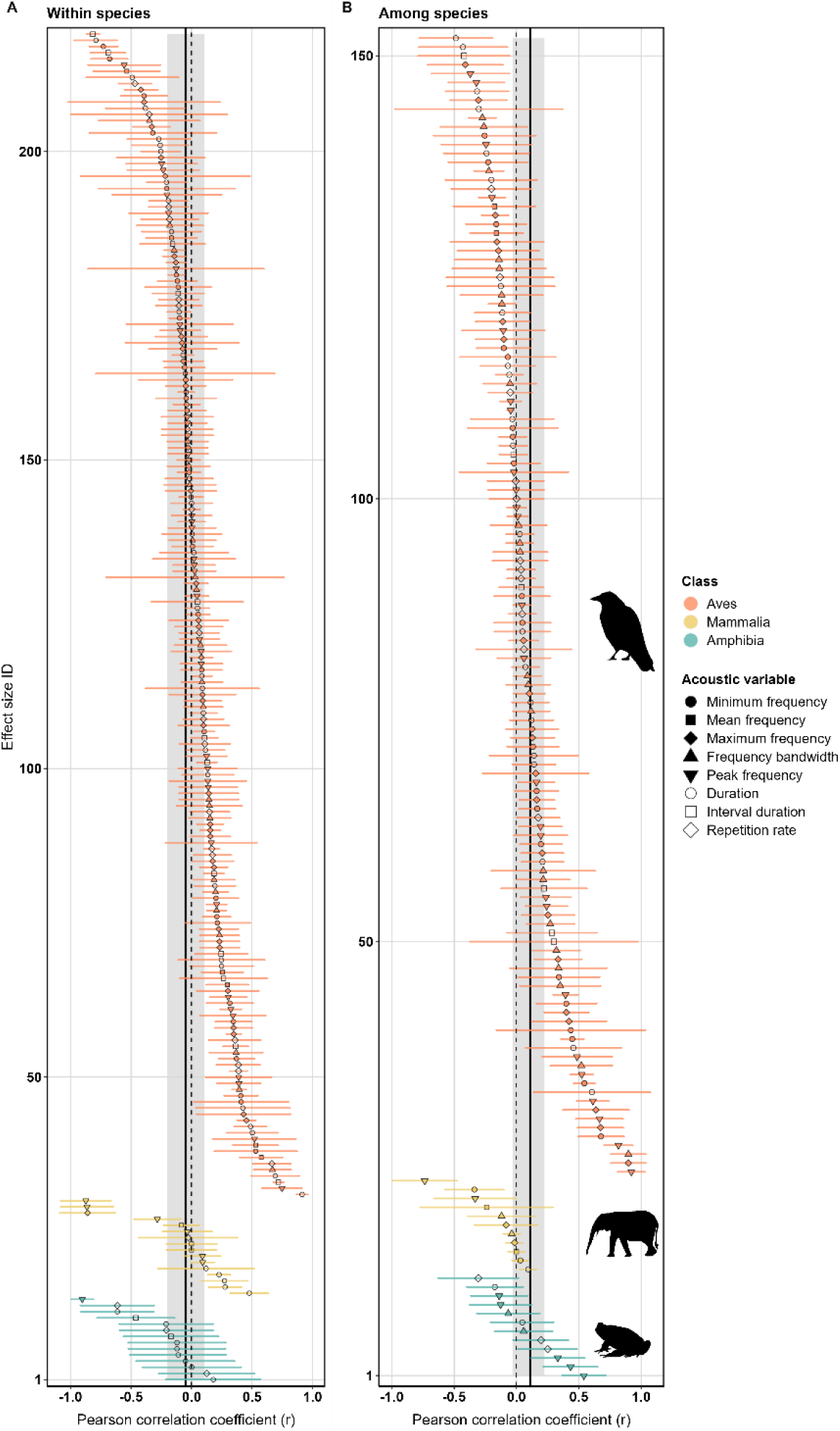
Caterpillar plot for (A) within- and (B) among-species comparisons. Effect sizes are displayed by taxonomic class, ordered by mean estimate, and with horizontal bars showing 95% confidence intervals (CIs). Symbols of the point estimate illustrate the acoustic variable measured. Positive Pearson correlation coefficients (*r*) indicate support for the acoustic adaptation hypothesis. Solid vertical black lines indicate the estimated global effect size obtained from the REML model with random factors only, and the grey area denotes the 95% CIs (A: *r* = -0.049, 95% CIs: -0.202 to 0.104; B: *r* = 0.111, 95% CIs: -0.031 to 0.254).

Our dataset included 22 studies (approximately 29% of our dataset) that were already included in a previous meta-analysis focusing on birds by Boncoraglio & Saino (2007). Nonetheless, we obtained different effect sizes for 10 (45 %) of these studies. Out of a total of 78 effect sizes reported in Boncoraglio & Saino (2007), 7 were identical with respect to their absolute value but had the opposite sign, 11 differed by more than 0.1, with 4 of them having the opposite sign, 2 effect sizes could not be retrieved, and 9 differed due to a different analytical approach (most probably because, contrary to Boncoraglio & Saino (2007), we did not pool data from different syllables or songs). Moreover, we excluded13 effect sizes (from 4 studies) used in the previous meta-analysis because the studies did not fit our selection criteria (i.e., habitats were impossible to classify as open or closed with regard to vegetation structure, habitat was not replicated, the study focused on the effects of urbanization, or the reported data were not sufficient to compute effect sizes). We report these inconsistencies in detail in Table S4.

### (2) Test of the AAH for within- and among-species comparisons

Overall, we did not find support for the AAH, neither for within-nor for among-species comparisons (Figs 3 and S3, Tables 1 and S6). None of the tested moderator variables explained a significant fraction of heterogeneity in effect sizes obtained from within- or among-species comparisons (Figs S4 and S5, and Tables S5 and S7).

**Fig. 3.**
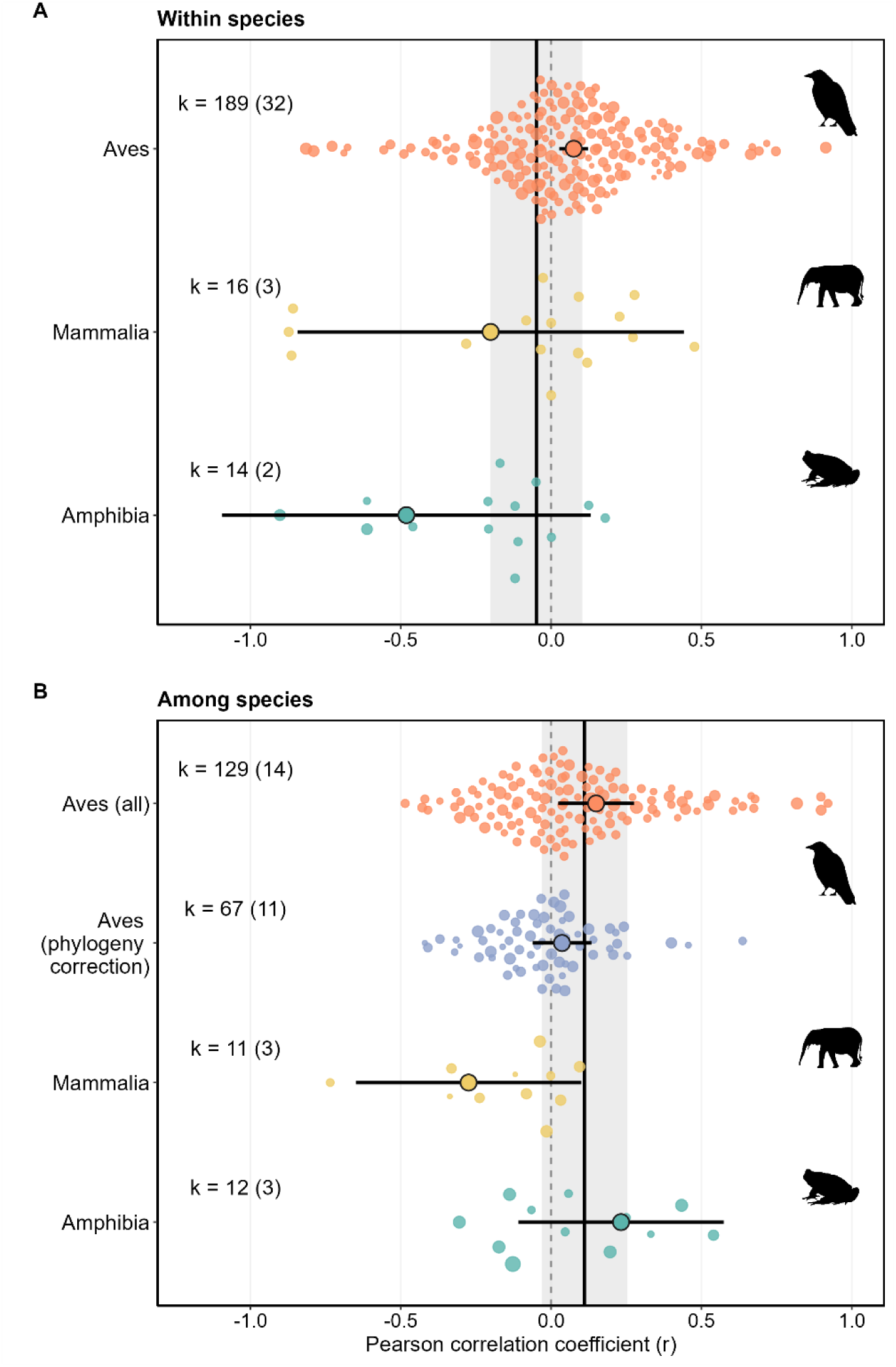
Orchard plots for (A) within- and (B) among-species comparisons showing effect sizes (Pearson correlation coefficients, *r*) and number of taxa (k = total number of effect sizes (total number of taxa) and mean effect size from REML models. Solid horizontal bars are 95% confidence intervals (CIs) of global effect sizes obtained for each taxon. Solid vertical black lines indicate the estimated global effect size from the REML model with the random factors only, and grey bands denote the 95% CIs (A: *r* = -0.049, 95% CIs: -0.202 to 0.104; B: *r* = 0.111, 95% CIs: -0.031 to 0.254). Circle size reflects precision of effect sizes (log(1/SE)).

When focusing only on the most studied taxonomic class, Aves, we found weak support for the AAH for within- and among-species comparisons (Figs 3 and S3). Frequency bandwidth showed a weak support for the AAH in within-species comparisons (Fig. S6), whereas minimum, maximum and peak frequencies showed significant support in among-species comparisons (Fig. S7). Analyses of other acoustic variables did not support the AAH. Primary studies of among-species comparisons obtained significantly higher effect sizes when they did not account for phylogenetic non-independence than when they did (Tables S5 and S7). When focusing on primary studies that accounted for phylogenetic non-independence, we did not obtain a significantly positive global effect size for birds, which was also the case when testing individual acoustic variables separately (Fig. S8). Results remained consistent when using different values of ρ (0.3, 0.7, and 0.9) to construct the variance-covariance matrix quantifying the non-zero covariance between sampling errors within the same primary studies (Table S8).

In analyses of mammals and amphibians, we found no support for the AAH neither in within-nor in among-species comparisons (Figs 3 and S3, Tables 1 and S6). For these analyses, the number of effect sizes and species was much lower compared to birds, which did not allow testing for the impact of moderators in these taxonomic classes.

The effect sizes we retrieved were highly heterogeneous (REML *I*² Total > 0.80 for all models; Table 1). For studies of within-species comparisons, heterogeneity arose mostly from variance among observations. For studies of among-species comparisons, heterogeneity stemmed primarily from among-study differences for the general model and the model including only birds. For the model including only studies on birds that accounted for phylogenetic non-independence (phylogeny correction), heterogeneity arose mostly from variance among species. In models including only mammals or amphibians, phylogeny explained the highest proportion of total heterogeneity (in both within- and among-species comparisons). These patterns were confirmed by the Bayesian approach, which generally estimated higher heterogeneity for among-study and among-observation differences compared to other factors, except for models with only mammals or amphibians (Table S6).

**Table 1.** Global effect sizes for the acoustic adaptation hypothesis obtained from the restricted maximum likelihood approach. Results of intercept-only General Linear-Mixed Effects Models are shown for the entire dataset (global model) and for subsets with respect to animal class (Aves, Mammalia or Amphibia). For among-species comparisons of birds, the results are shown for a subset of studies that accounted for phylogenetic non-independence (Aves (phylogeny correction)). Table shows number of effect sizes (k), number of species or taxonomic groups (N), estimates of *r* with 95% confidence limits in parentheses and heterogeneity *I*² (total, variance arising from phylogenetic affinities, taxon, study identity, and observation).

| Type of study | Taxonomic range | k | N | *r* |  | z-value | P-value | *I*<sup>2</sup> Total | *I*<sup>2</sup> Phylogeny | *I*<sup>2</sup> Taxon | *I*<sup>2</sup> Study | *I*<sup>2</sup> Observation |
| --- | --- | --- | --- | --- | --- | --- | --- | --- | --- | --- | --- | --- |
| Within species | Global model | 219 | 37 | -0.049 | (-0.202,0.104) | -0.622 | 0.534 | 0.955 | 0.138 | 0.000 | 0.280 | 0.537 |
|  | Aves (all) | 189 | 32 | 0.075 | (0.027,0.123) | 3.057 | **0.002** | 0.937 | 0.000 | 0.000 | 0.000 | 0.937 |
|  | Mammalia | 16 | 3 | -0.201 | (-0.844,0.441) | -0.614 | 0.539 | 0.979 | 0.000 | 0.886 | 0.000 | 0.092 |
|  | Amphibia | 14 | 2 | -0.482 | (-1.096,0.132) | -1.537 | 0.124 | 0.892 | 0.272 | 0.272 | 0.272 | 0.076 |
| Among species | Global model | 152 | 20 | 0.111 | (-0.031,0.254) | 1.532 | 0.126 | 0.950 | 0.055 | 0.025 | 0.642 | 0.228 |
|  | Aves (all) | 129 | 14 | 0.150 | (0.024,0.276) | 2.331 | **0.020** | 0.945 | 0.000 | 0.020 | 0.692 | 0.233 |
|  | Aves (phylogeny<br>correction) | 67 | 11 | 0.036 | (-0.062,0.134) | 0.724 | 0.469 | 0.820 | 0.000 | 0.385 | 0.000 | 0.435 |
|  | Mammalia | 11 | 3 | -0.275 | (-0.649,0.100) | -1.435 | 0.151 | 0.972 | 0.611 | 0.000 | 0.328 | 0.033 |
|  | Amphibia | 12 | 3 | 0.232 | (-0.109,0.574) | 1.334 | 0.182 | 0.865 | 0.474 | 0.000 | 0.000 | 0.390 |

### (3) Sensitivity analyses and publication bias

Sensitivity analyses revealed that model estimates are robust and reliable. Outlier analyses for estimates of *r* of the complete datasets of within- and among-species comparisons (Grubb’s test of *r:* G = 3.045, P-value = 0.230; G = 2.922, P-value = 0.234; respectively, Fig S9). This was also the case for within- and among-species comparisons of birds (G = 3.075, P-value = 0.176; G = 2.86, P-value = 0.236; respectively, Fig. S9). An outlier was detected when focusing on primary studies that accounted for phylogenetic non-independence (i.e., maximum frequency estimate for Parulidae, Grubb’s test of *r* G = 3.390, P-value = 0.013). We found that there were no influential studies having disproportionate effects on the model estimates (Fig. S10). Leave-one-out analysis demonstrated that removing one study at a time did not change the results of within-species comparisons for both the global model and the model including only birds. In among-species comparisons, the removal of four studies (one on birds and three on mammals) resulted in marginally overall positive support for the AAH (Fig. S10 B). However, when focusing solely on studies on birds, the removal of the study on birds did not affect the overall results (Fig. S10 D). Using Fisher’s *z* instead of the Pearson correlation coefficient as an effect size provided very similar results suggesting our analysis to be robust with respect to the chosen effect size metric (Table S9).

We detected no evidence for the small-study effect either for within- and among-species comparisons, since small studies (with smaller sample size and lower precision) did not show stronger effect sizes than large studies (Figs S11A and B). In fact, for studies reporting within-species comparisons, we detected the opposite trend, i.e., a significant negative relationship between effect size and its standard error (Fig. S11A). In addition, we found no evidence for funnel plot asymmetry (Figs S11C, D, E, and F) inferred from Kendall’s rank correlation test (Kendall’s tau [within-species comparisons] = -0.059, *P-value* = 0.212; Kendall’s tau [among-species comparisons] = -0.005, *P-value* = 0.938).

Finally, we also detected no evidence for a time-lag bias since the year of publication did not predict effect sizes of within-species comparisons (slope = 0.001, 95% CI = [-0.006, 0.009], z-value = 0.380, P-value = 0.705) or among-species comparisons (slope = -0.007, 95% CI = [-0.016, 0.001], z-value = -1.791, P-value = 0.073). We observed a negative relationship for studies not accounting for phylogenetic non-independence and a slightly positive relationship for studies that accounted for it, but these relationships were not statistically significant (Figs S11G and H).

## IV. DISCUSSION

The acoustic adaptation hypothesis (AAH) posits that animals produce acoustic signals that have evolved in response to the structure of their habitat, but ever since its original formulation, there has been much uncertainty about the hypothesis’ validity and generality within and among species. We analysed a dataset of 371 effect sizes across 75 studies and including birds, mammals, and amphibians to demonstrate that acoustic variables do not differ significantly between habitats with different vegetation structure in both within- and among-species comparisons. Therefore, current empirical evidence does not support the idea that the AAH provides a general explanation for the evolution of acoustic signals. Only within-species comparisons of birds provided weak support for this hypothesis indicating that variation in acoustic signals in relation to vegetation structure may be more complex and context-dependent than previously thought. Below, we discuss these findings, identify the limitations of our study, and propose directions for future research.

### (1) Limited support for the AAH

Our phylogenetically informed synthesis provides no overall support for the AAH, offering quantitative evidence that acoustic signals do not show predictable responses to vegetation structure across vertebrates. This result might seem unexpected given the established principles of the physics of sound transmission (Hardt & Benedict, 2021) and questions the broader utility of the AAH for understanding the evolution of acoustic communication in natural settings.

We acknowledge that our final dataset exhibits a taxonomic bias with an overrepresentation of birds. While this reflects a limitation of our study, it also underscores the evident need for additional research examining the influence of vegetation structure on acoustic signals of taxa beyond birds—a call that has been made by numerous authors before (e.g., Ey & Fischer, 2009; Goutte *et al*., 2018; Mikula *et al*., 2021) but remains largely unaddressed. As research extends to taxa that are currently understudied, the inclusion of reptiles, insects, and more species of anurans and mammals would likely enable the examination of other potential moderators, such as signal development (i.e., innate versus learned acoustic signals), which has been suggested suggested by other authors to influence results, but that could not be tested in this study (see Text S1). This would ultimately contribute to a more comprehensive understanding of acoustic adaptation and animal communication.

The substantial variation in effect sizes across our dataset (REML *I*² Total > 0.80 for all the models; Table 1), a common pattern of meta-analyses in ecology and evolution (with the average *I*^2^ being as high as 92%; Senior *et al*., 2016), remained largely unexplained. For within-species comparisons, most of the heterogeneity originated from variation among observations, while differences among studies, taxa or phylogeny effects had less impact. This among-observation variability may reflect the contrasting support of the AAH within studies depending on the acoustic variables analysed, sometimes even within the same acoustic variables when measured for the entire signal and its fragments (i.e., for studies that measured acoustic variables across the entire song as well as its syllables). On the other hand, the heterogeneity of among-species comparisons arose mostly from variation among studies, which could be due to differences in study design such as using a different assessment of vegetation structure, a focus on different acoustic variables, or differences in statistical correction for body size or phylogenetic non-independence. Remarkably, only a fraction of primary studies of among-species comparisons accounted for phylogenetic non-independence and our analyses suggest that this methodological aspect has an impact on the study’s outcome, particularly in birds. In contrast to a qualitative review by Ey & Fischer (2009), our results suggest that studies not accounting for phylogenetic non-independence tend to obtain stronger support for the AAH than those that do. This underscores the need for comparative studies to consider phylogenetic relatedness to avoid misinterpreting the role of vegetation structure in shaping acoustic signals.

Why then does the AAH not seem to apply universally? Several factors may contribute to the limited support for the AAH. First, the complexity of the habitat and methodological difficulties regarding its classification could render the AAH an oversimplified concept if applied too broadly. Second, the diversity of acoustic signals across taxa and the multiplicity of selective pressures acting on them may lead to complex and diverse responses that may override a potential effect of the habitat.

#### (a) Complexity of the habitat

Although the AAH predicts that selection should favour acoustic signals that propagate well in an environment, it has been primarily tested in terms of effects of vegetation on sound transmission. This approach might be too simplistic, as many other elements may influence sound propagation in a given habitat (Attenborough & Renterghem, 2021). Physical features of the landscape (hills, valleys, or slopes), type of substrate, weather conditions, as well as background noise (arising from wind, moving water, or other species) are only a few of the factors that can also affect how sound travels (Sueur, Krause & Farina, 2019; Guibard *et al*., 2022). For example, there is increasing evidence that animals can adjust their acoustic signals to surrounding noise, with most of the studies highlighting within-species adjustments (Slabbekoorn & Peet, 2003; Gil *et al*., 2015; Gomes *et al*., 2022). The significant heterogeneity detected in our analyses probably reflects variation among studies arising not only from methodological aspects including differences in data collection, analysis, and reporting but also from the physical features of the habitat.

Even when focusing on vegetation structure only, characterizing the habitat for the purpose of investigating animal acoustic communication proves to be a challenging and not yet standardized task. Early work on the AAH used broad habitat categories (forest, edge, and grassland; Morton, 1975) and general predictions were then formulated for open versus closed habitats. These classifications of vegetation structure do not only oversimplify specific habitat characteristics crucial for sound transmission but also tend to be subjective and prone to ambiguity. Even the use of more quantitative continuous variables does not resolve this issue because a wide array of both small-scale (measured at a microhabitat level, e.g., number of trees) and large-scale (representative of macrohabitat characteristics, e.g., tree cover, canopy cover) variables have been used, which challenges the generalization of findings. The differences in the assessment of vegetation structure among studies may have contributed to the heterogeneity in our results as suggested by Ey & Fischer (2009). However, we found that support for the AAH did not significantly differ among studies using categorical versus continuous characterization of vegetation structure, or among studies with varying magnitudes of contrast between closed and open habitats. This indicates that including different classifications of the habitat did not change the overall results, possibly reflecting the lack of power of the current classification to capture vegetation structure. Additionally, within the same habitat, both season and the time of day can have a significant influence on signal transmission. For example, an empirical study conducted in two different forests demonstrated that these factors impacted detection distance by up to a factor of five (Haupert, Sèbe & Sueur, 2022).

#### (b) Alternative determinants of signal evolution overriding habitat effects

Animal acoustic signals are shaped by a multitude of selective forces, which can give rise to contrasting responses, potentially reducing the significance of habitat effects, thus challenging the universality of the AAH. Whilst our results provide no overall evidence for the AAH, we found weak support for the AAH in birds for within-species comparisons. Avian acoustic signals might therefore present phenotypically plastic responses to vegetation structure but the weak support observed in our analysis suggests a minor role, if any, of the vegetation in shaping acoustic signals even in birds. The lack of support for the AAH in mammals and amphibians aligns with the previous qualitative review (Ey & Fischer, 2009) and may reflect the limited importance of habitat in explaining global signal variation in these taxa. Nevertheless, these results demand caution due to the limited number of species tested for these animal classes.

Further complicating the application of the AAH as a universal framework are diverse taxon-specific constraints associated with each acoustic variable. Anuran calls, for instance, exhibit greater morphological constraints on frequency than on temporal parameters (Escalona Sulbarán *et al*., 2019; Tonini *et al*., 2020), and this has been suggested as the reason for temporal parameters being more susceptible to evolve in response to environmental factors compared to spectral ones (Mendoza-Henao *et al*., 2023). For birds, there seems to be no consensus regarding the acoustic parameters that are more prone to be selected by environmental factors. While the previous meta-analysis found that all frequency variables differed between habitats, consistently supporting the AAH (Boncoraglio & Saino, 2007), Ey & Fischer (2009) identified minimum frequency and frequency bandwidth as the only acoustic variables presenting global support for the AAH. Recent findings propose frequency bandwidth as a more labile acoustic variable, in opposition to peak frequency, which is more constrained by a species’ genetic background (Rivera *et al*., 2023). Although our results focussing on plastic changes (i.e. within-species comparisons) align with the findings from Rivera *et al*. (2023), this is not the case for the results obtained for evolutionary changes (i.e. among-species comparisons), which reveal a lack of support for all the acoustic variables when using studies that accounted for phylogeny.

Even if animals do not adjust acoustic parameters to vegetation structure, they can exhibit other adaptive communication strategies to optimize signal transmission. In particular, they may use aerial displays (Menezes & Santos, 2020) or move to a more suitable microhabitat and height for broadcasting (Kime, 2000; Seddon, 2005; Chitnis, Rajan & Krishnan, 2020) or for listening (Jensen, Larsen & Attenborough, 2008). In the presence of background noise from biological, abiotic, or urban sources, animals may also adjust the broadcasting time to minimize signal overlap and competition for ’signal space’ with other sound-producing species (Hart *et al*., 2015; Chronister, Rhinehart & Kitzes, 2023; Staniewicz *et al*., 2023). Ultimately, the benefits of long-distance transmission, such as improved communication over larger distances or enhanced mate attraction, may be offset by the associated predatory, competitive, and parasitic costs (Ryan *et al*., 1981; Garamszegi & Avilés, 2005).

### (2) Changing the paradigm

As Brown & Handford (1996) concluded: “*With all the possible selective forces acting on signal design, investigators looking for specific or finely detailed differences within or among species, across habitats, should not be surprised to find results that do not always agree with AAH predictions*”. This conclusion is echoed in more recent work (e.g., Ey & Fischer, 2009; Goutte *et al*., 2018; Mikula *et al*., 2021) but the AAH has remained a focal point of investigation.

The lack of support for the AAH in our analyses calls for a more comprehensive and integrative approach to study acoustic adaptation, considering the complex interplay of ecological and evolutionary factors. We propose this field to move beyond the current confines of the AAH by highlighting promising avenues for future studies.

#### (a) Standardizing habitat attenuation measurements to refine AAH predictions

We believe that advances in understanding how environmental features affect acoustic signalling can be achieved through a different approach of how the habitat is characterised and analysed. This involves refining the description of habitats and adopting a broader perspective, moving beyond a sole focus on vegetation structure. Ey & Fischer (2009) suggested a detailed characterization of each habitat, including altitude, humidity, temperature, local animal acoustic signals, and vegetation structure. However, obtaining and interpreting such an extensive amount of information at the relevant scale for communication may pose practical challenges due to the multitude of factors involved and difficulties in comparing studies. Instead of considering multiple factors of the habitat separately, it may be preferable to use a single variable that integrates the direct effects of multiple factors on sound propagation.

A promising approach to characterise the acoustic properties of the environment has been emphasised recently by Haupert *et al*. (2022). By recording ambient sound and broadcasting white noise and recording it at various distances from the emitter in two forest types, they found that the coefficient of attenuation (a_0_, expressed in dB/kHz/m) can be used to describe accurately the attenuation characteristic of an environment for large frequency bandwidths (for 5 kHz frequency bandwidths between 0-20 kHz in their study). This singular parameter is modelled based on acoustic physics rather than descriptive statistics and summarizes the contribution of various variables, including temperature, relative humidity, atmospheric pressure, distance, frequency, ambient sound pressure level, and vegetation, to global attenuation (Haupert *et al*., 2022). Moreover, Haupert *et al*. (2022) used the same playback technique to describe the attenuation spectrum (the pattern of attenuation by frequency) of the two studied forests. Despite having similar a_0_ values, both forests exhibited very different attenuation spectrum profiles. Within the same environment, detection distance varied by up to a factor of five depending on the frequency considered. These spectrum profiles potentially reveal frequencies with the highest detection probability and could be used as starting point to describe the habitat. Subsequently, studies could establish correlations between the attenuation profile and the species’ (or population’s, or even individual’s) specific power spectrum of acoustic signals, in order to test whether frequencies with greater detection distance are preferentially used. New research should also consider that within the same signal, for example a song, there might be a maximum transmission distance for the detection (i.e., the hearing and perception of a conspecific through its song) and for decoding of the signal (i.e., deciphering the conspecific’s performance from its song). In summary, we suggest a more comprehensive examination of attenuation in each locality, employing methodologies such as those outlined by Haupert *et al*. (2022). These methodologies are applicable in any environment and encompass all variables that might influence sound, thus offering a more holistic approach. This would facilitate global comparisons (through a single variable) and offer a more objective and standardized assessment of the acoustic properties of a given habitat.

Finally, we propose to rephrase the predictions of the AAH to take into account the attenuation spectrum. Specifically, animals inhabiting environments with different attenuation spectra are predicted to emit acoustic signals at frequencies (minimum, mean, maximum, and peak frequency) and times (duration, interval duration, repetition rate) such that detection and/or decoding distances are maximized. This implies that there is no universal pattern for the expected change in these variables, unless habitats share similar profiles. Instead, the alterations will be contingent upon the specific characteristics of the environment. This adjustment also overcomes an earlier noticed limitation of the AAH that for maximum efficiency, long-range acoustic communication in any habitat should employ the lowest possible frequencies (Wiley & Richards, 1982).

#### (b) Towards a narrower framework to test the AAH

While the AAH offers valuable insights, it may not fully capture the processes underlying the evolution of acoustic signalling as it does not account for alternative strategies that animals use in response to multiple selective pressures acting on signal structure or their interactions. To comprehensively assess the environmental influence on signal evolution, a more comprehensive approach is necessary.

Considering the detected lack of generality of the AAH, it would be valuable to narrow down the focus to signals (or specific parts of signals) for which an advantage in transmission over long distances is likely. Studies not specifically selecting for these signals (e.g., Mikula *et al*., 2021) might reflect this limitation. For example, many acoustic signals are part of private duets (i.e., do not need long-distance transmission) such that there is no selection against easily degrading communication (Larsen, 2020). Likewise, for some species song is not used for distant communication, as reported in zebra finches, where it may be more pertinent to assess whether their distance call, rather than their song, is adapted for long distance transmission through the local environment (Loning, Griffith & Naguib, 2022). This approach requires an in-depth knowledge of the studied species, which might not always be readily available and can be challenging to obtain. Nevertheless, we believe that tests of the AAH are only meaningful for signals that are meant for long-distance communication.

## V. CONCLUSIONS

1. Our meta-analysis challenges the general validity of the acoustic adaptation hypothesis (AAH) by showing no overall support for the correlation between vegetation structure and acoustic variables.
2. The weak support for the AAH in within-species comparisons of birds suggests a phenotypically plastic response of frequency bandwidth to the vegetation structure of the habitat. By contrast, the AAH is not supported by among-species comparisons of studies accounting for phylogenetic non-independence.
3. Although many years have passed since the formulation of the AAH, major efforts are still required to confirm the predictive ability of the hypothesis. Future research should refine the characterization of the habitat through a standardized quantification of the attenuation, which will lead to habitat- and species-specific predictions. Moreover, we expect that major advances in the field will benefit from broadening the studied taxonomic range while concurrently being selective about the tested signals.

## ACKNOWLEDGEMENTS

We are very grateful to the authors of the primary studies for making their research available, and in particular those providing additional data or information, and unpublished datasets, including Emma Greig, Elodie Ey, Darren Proppe, and Martim Melo. T. Michael Keesey, Julián Bayona, and Dennis Murphy designed the silhouettes used in our figures. We thank Sylvain Haupert, Ole Næsbye Larsen, Frédéric Sèbe, and Keith Attenborough for their insightful comments and discussions, which significantly improved the manuscript. We extend our gratitude to Rémi Tournebize for his assistance with troubleshooting R scripts.

Funding: B.F. was funded by the Foundation for Science and Technology (FCT, Portugal) through a PhD grant (2020.04569.BD). This research was partly funded by grant PID2021-124501NB-I00 to B.M. from the Spanish Ministry of Science and Innovation, co-financed by the European Union’s Regional Development Fund (ERDF). C.T. benefited from a grant managed by the Agence Nationale de la Recherche (CEBA: ANR-10-LABX-25-01). T.J. received funds from the Centre National de la Recherche Scientifique (CNRS).

## VI. AUTHOR CONTRIBUTIONS

B.F. Conceptualization, Data curation, Formal analysis, Investigation, Methodology, Project administration, Visualization, Writing – original draft;

P.B.D. Supervision, Writing – review and editing

B.M. Supervision, Writing – review and editing

C.T. Supervision, Writing – review and editing

T.J. Conceptualization, Data curation, Formal analysis, Investigation, Methodology, Project administration, Supervision, Validation, Visualization, Writing – review and editing

## VII. DATA AVAILABILITY

All data and R scripts associated with this study can be retrieved from the OSF data repository (it will be made public upon article acceptance).

## X. SUPPORTING INFORMATION

Additional supporting information may be found online in the Supporting Information section at the end of the article.

**Text S1.** Differences between the pre-registered methods and the ones used are explained in this document.

**Table S1.** Definition of each acoustic variable and the general predictions of the acoustic adaptation hypothesis for a closed habitat in comparison to an open habitat.

**Table S2.** Source data table of effect sizes included in the meta-analysis.

**Table S3.** Number of studies and effect sizes for each moderator category included in this meta-analysis for primary studies of within- and among-species comparisons.

**Table S4.** Studies and effect sizes included in a previous meta-analysis testing the acoustic adaptation hypothesis in birds by Boncoraglio & Saino (2007) and effect sizes included in our study converted to Fisher’s z. Inconsistencies between this and our study were divided in six different categories: (1) effect size differs more than 0.10; (2) effect size is similar but differs in sign; (3) effect size could not be retrieved for our study; (4) study excluded from our meta-analysis based on our selection criteria; (5) different methodological approach: authors might have pooled the data and we did not; (6) different methodological approach: we used raw values.

**Table S5.** Summary statistics of moderator models of within- and among-species comparisons using restricted maximum likelihood approach. Moderators include habitat classification (categorical versus continuous), habitat contrast of categorical classifications (0 versus 1), and body size correction (no versus yes). For comparisons among species, moderators also include phylogeny correction (no versus yes). The first category mentioned in parentheses is used as the reference level for the model results. Phylogenetically controlled multilevel meta-analytic single predictor models are shown with Omnibus tests (Q_M_, Wald-type chi-square test) and pseudo-R².

**Table S6.** Global tests using the Bayesian approach. Results of intercept-only general linear-mixed effects models are shown for the entire dataset (global model) and subsets with respect to animal class (Aves, Mammalia or Amphibia). For among-species comparisons of birds, the results are also shown for a subset of studies that accounted for phylogenetic non-independence (‘Aves (phylogeny correction)’). Table shows number of effect sizes (k), number of species or taxonomic groups (N), estimates of *r* with 95% confidence intervals in parentheses and heterogeneity *I*² (variance arising from phylogenetic affinities, taxon, study identity, and observation). Model estimates are shown as posterior modes with 95% Highest Posterior Density (HPD) intervals in brackets.

**Table S7.** Test of moderators explaining for within- and among-species comparisons using the Bayesian approach. Moderators include habitat classification (categorical versus continuous), habitat contrast of categorical classifications (0 versus 1), and body size correction (no versus yes). For comparisons among species, moderators also include phylogeny correction (no versus yes). The first category mentioned in parentheses is used as the reference level for the model results. Model estimates (i.e., estimated difference between groups) are shown as posterior modes with 95% Highest Posterior Density (HPD) intervals obtained from phylogenetically controlled general linear-mixed effects models. The variance explained by the entire model and by the fixed factors is given as the conditional R² and the marginal R², respectively, with 95% HPD intervals in brackets.

**Table S8.** Sensitivity analysis examining the robustness of the overall effect size (r) to the assumption of different ρ values (within-study correlation) used to construct the variance-covariance metrics that account for non-independence of sampling variances. Table shows estimates of r with 95% confidence intervals in parentheses.

**Table S9.** Global effect sizes (Fisher’s z) for the acoustic adaptation hypothesis obtained from the restricted maximum likelihood approach. Results of intercept-only General Linear-Mixed Effects Models are shown for the entire dataset (global model) and for subsets with respect to animal class (Aves). For among-species comparisons of birds, the results are shown for a subset of studies that accounted for phylogenetic non-independence (Aves (phylogeny correction)). Table shows number of effect sizes (k), number of species or taxa (N), and estimates of Fisher’s z and the 95% confidence intervals in parentheses.

**Fig. S1.** Phylogeny used to account for phylogenetic non-independence in statistical analysis of within-species comparisons. Doughnut charts show the proportion of sample sizes (i.e., number of Pearson correlation coefficients) and the number of species.

**Fig. S2.** Phylogeny used to account for phylogenetic non-independence in analysis of among-species comparisons. Doughnut charts show the proportion of sample sizes (i.e., number of Pearson correlation coefficients) and the number of taxa.

**Fig. S3.** Density plots showing posterior distributions obtained from Bayesian models for the differences in Pearson correlation coefficients (r) among animal classes obtained from general linear mixed-effects models (see Methods). Positive values indicate support for the AAH.

**Fig. S4.** Orchard plots for within-species comparisons showing individual effect sizes and mean effect size obtained from REML models for each acoustic variable. Number of effect sizes and number of taxa (k = number of effect sizes (number of taxa)) are shown on the left for each variable. Bold horizontal bars are 95% confidence intervals (CIs) and thin black lines are precision intervals. Positive Pearson correlation coefficients (r) indicate support for the acoustic adaptation hypothesis. Solid vertical line indicates the estimated global effect size obtained from the REML model with random factors only, and the grey area denotes the 95% CIs (*r* = -0.049, 95% CIs: -0.202 to 0.104). Circle size reflects effect size precision (log(1/SE)).

**Fig. S5.** Orchard plots for among-species comparisons showing individual effect sizes and mean effect size from REML models for each acoustic variable. Number of effect sizes and number of taxa (k = number of effect sizes (number of taxa)) are shown on the left for each variable. Bold horizontal bars are 95% confidence intervals (CIs) and thin black lines are precision intervals. Positive Pearson correlation coefficients (r) indicate support for the acoustic adaptation hypothesis. Solid vertical line indicates the estimated global effect size obtained from the REML model with random factors only, and the grey area denotes the 95% CIs (*r* = 0.111, 95% CIs: -0.031 to 0.254). Circle size reflects effect size precision (log(1/SE)).

**Fig. S6.** Orchard plots for comparisons within bird species showing individual effect sizes and mean effect size obtained from REML models for each acoustic variable. Number of effect sizes and number of taxa (k = number of effect sizes (number of taxa)) are shown on the left for each variable. Bold horizontal bars are 95% confidence intervals (CIs) and thin black lines are precision intervals. Positive Pearson correlation coefficients (r) indicate support for the acoustic adaptation hypothesis. Solid vertical line indicates the estimated global effect size obtained from the REML model with random factors only, and the grey area denotes the 95% CIs (*r* = 0.075, 95% CIs: 0.027 to 0.123). Circle size reflects effect size precision (log(1/SE)).

**Fig. S7.** Orchard plots for comparisons among bird species showing individual effect sizes and mean effect size from REML models for each acoustic variable. Number of effect sizes and number of taxa (k = number of effect sizes (number of taxa)) are shown on the left for each variable. Bold horizontal bars are 95% confidence intervals (CIs) and thin black lines are precision intervals. Positive Pearson correlation coefficients (r) indicate support for the acoustic adaptation hypothesis. Solid vertical line indicates the estimated global effect size obtained from the REML model with random factors only and the grey area denotes the 95% CIs (*r* = 0.150, 95% CIs: 0.024 to 0.276). Circle size reflects effect size precision (log(1/SE)).

**Fig. S8.** Orchard plots for comparisons that accounted for phylogenetic non-independence (phylogeny correction) among bird species showing individual effect sizes and mean effect size from REML models for each acoustic variable. Number of effect sizes and number of taxa (k = number of effect sizes (number of taxa)) are shown on the left for each variable. Bold horizontal bars are 95% confidence intervals (CIs) and thin black lines are precision intervals. Positive Pearson correlation coefficients (r) indicate support for the acoustic adaptation hypothesis. Solid vertical line indicates the estimated global effect size from the REML model including only phylogeny corrected studies with random factors only, and the grey area denotes the 95% CIs (*r* = 0.036, 95% CIs: -0.062 to 0.134). Circle size reflects effect size precision (log(1/SE)).

**Fig. S9.** Bar and density plots for within- (left) and among-species (right) comparisons showing the distribution of effect sizes (Pearson correlation coefficient (r)). Plots are shown for entire dataset (A and B) and for a subset of studies focusing on birds (C and D). Positive Pearson correlation coefficients (*r*) indicate support for the acoustic adaptation hypothesis. Solid vertical black lines indicate the estimated global effect size obtained from the REML model with random factors only for global models (A: *r* = -0.049; B: *r* = 0.111) and for the ones including only comparisons of birds (C: *r* = 0.075; D: *r* = 0.150).

**Fig. S10.** Caterpillar plots of leave-one-out analyses for within- (left) and among-species (right) comparisons based on datasets with one study left out at a time from model fitting. Plots are shown for entire dataset (A and B) and for a subset of studies focusing on birds (C and D). Estimated global effect sizes are ordered by mean estimate, and with horizontal bars showing 95% confidence intervals (CIs). Positive Pearson correlation coefficients (*r*) indicate support for the acoustic adaptation hypothesis. Solid vertical black lines indicate the estimated global effect size obtained from the REML model with random factors only, and the grey area denotes the 95% CIs for global models (A: *r* = -0.049, 95% CIs: -0.202 to 0.104; B: *r* = 0.111, 95% CIs: -0.031 to 0.254) and for the ones including only comparisons of birds (C: *r* = 0.075, 95% CIs: 0.027 to 0.123; D: *r* = 0.150, 95% CIs: 0.024 to 0.276).

**Fig. S11.** Plots for the assessment of publication bias for within- (left) and among-species (right) comparisons. Regressions (A and B) show the relationship between effect size (Fisher’s z) and its standard error testing whether studies with smaller sample sizes and lower precision are more likely to be published when reporting larger effect sizes. Solid lines show the fit of the multivariate linear mixed-effects model with year and all the moderators as fixed effects and study identifier, effect size identifier, taxon name, and phylogeny as random terms (within-species comparisons: slope = -0.887, 95% CI = [-1.766, -0.008], t-value = -1.990, P-value = 0.048; among-species comparisons: slope = - 0.206, 95% CI = [-1.382, 0.970], t-value = -0.346, P-value = 0.730). Circle size reflects precision of effect sizes (log(1/SE)). Funnel plots for raw values (C and D) and for meta-analytic residuals (E and F) were obtained from multivariate linear mixed-effects model with all the moderators as fixed effects and study identifier, effect size identifier, taxon name, and phylogeny as random terms. Dashed lines indicate the estimated global effect size. Bold and thin black solid lines denote the expected 95% and 99% confidence limits, respectively, purely due to sampling heterogeneity. Asymmetries along the global effect size may reflect publication biases. Time-lag bias analysis (G and H) showing the relationship (linear regression) between effect size (Pearson correlation coefficient, r) and the year of publication of studies that did (and did not) account for phylogenetic non-independence (within-species comparisons: slope ± SE = -0.001 ± 0.002, t-value = -0.545, P-value = 0.586; among-species comparisons: slope[yes] ± SE = 0.002 ± 0.004, t-value = 0.514, P-value = 0.609, slope[no] ± SE = - 0.005 ± 0.003, t-value = -1.625, P-value = 0.108). Circle size reflects effect size precision (1/SE).

## Notes

### Competing Interest Statement

The authors have declared no competing interest.

### Summary of Updates

Added sensitivity analyses and improved the text, figures and tables.

