## Supplementary Information for "Meta-analysis of the acoustic adaptation hypothesis reveals no support for the effect of vegetation structure on acoustic signalling across terrestrial vertebrates"

Author ORCIDs

Bárbara Freitas http://orcid.org/0000-0001-6020-6236

Pietro B. D'Amelio http://orcid.org/0000-0002-4095-6088

Borja Milá http://orcid.org/0000-0002-6446-0079

Christophe Thébaud http://orcid.org/0000-0002-8586-1234

Tim Janicke http://orcid. org/0000-0002-1453-6813

**SUPPORTING INFORMATION**

**FIGURES**

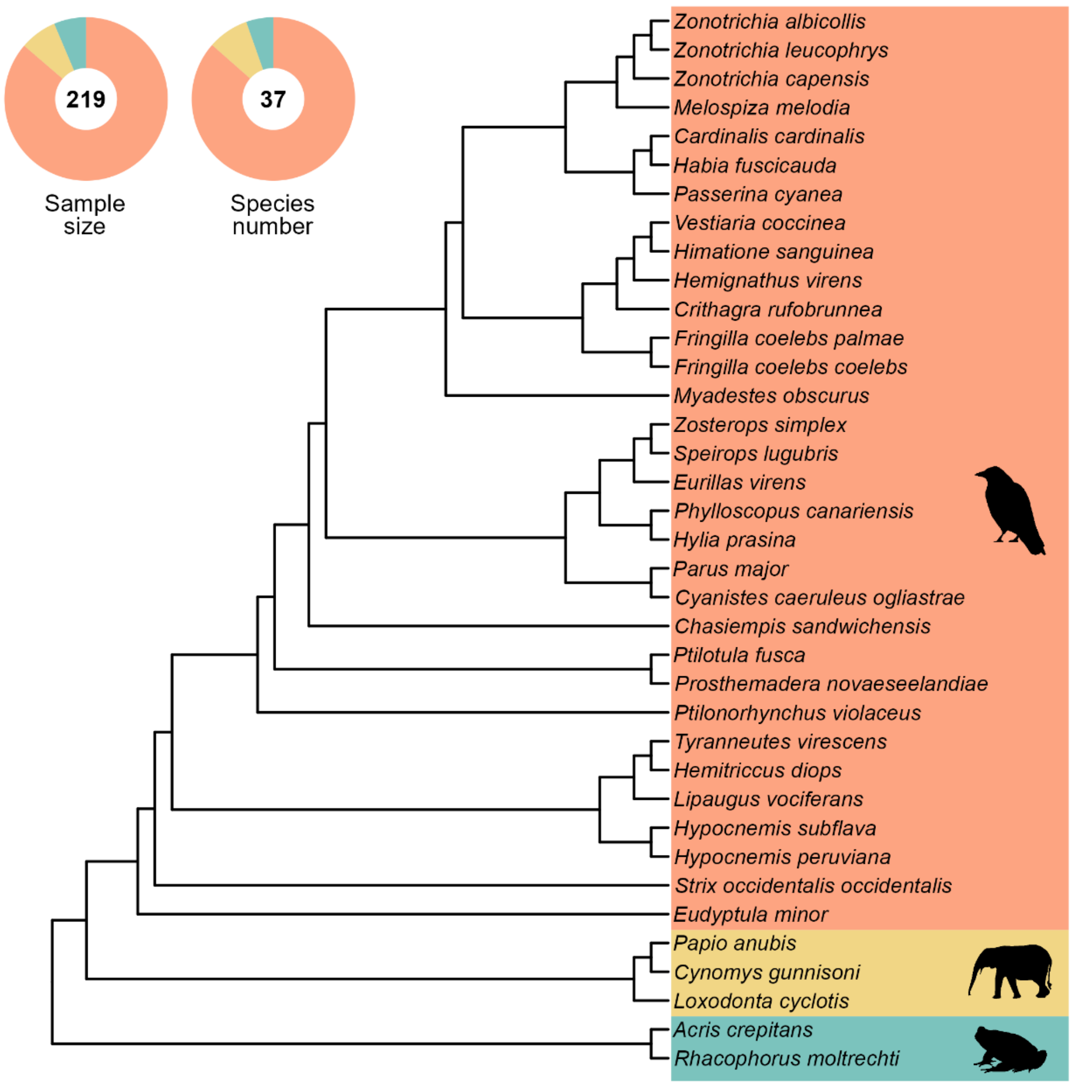


**Fig. S1.** Phylogeny used to account for phylogenetic non-independence in statistical analysis of within-species comparisons. Doughnut charts show the proportion of sample sizes (i.e., number of Pearson correlation coefficients) and the number of species.

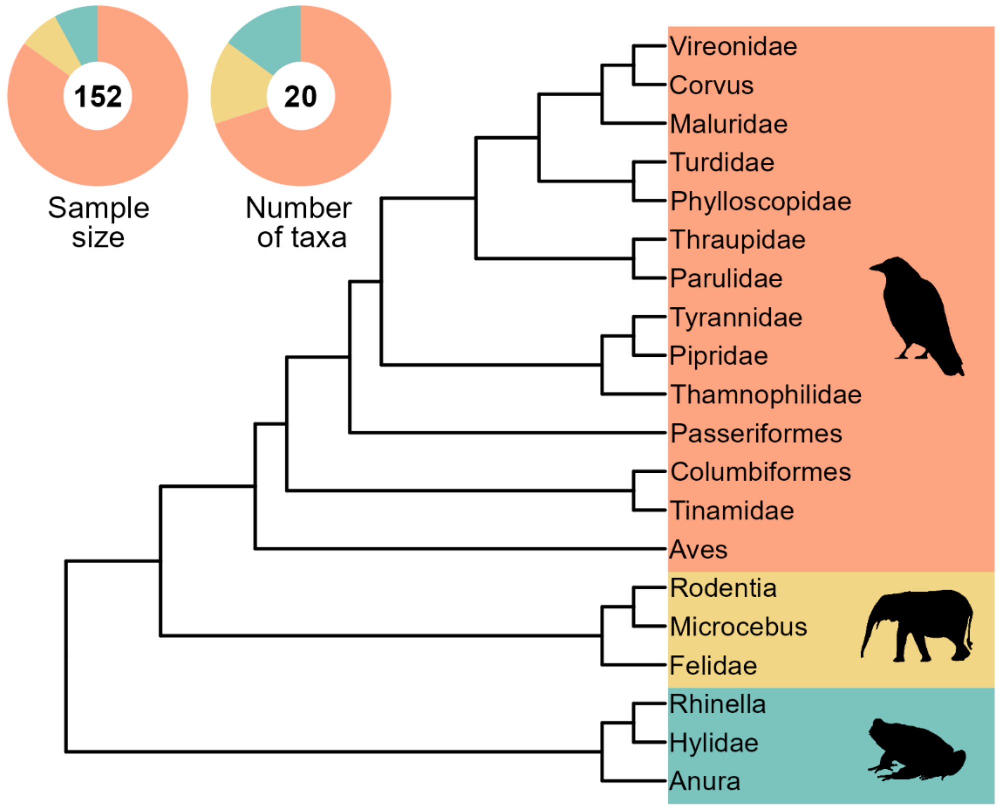


**Fig. S2.** Phylogeny used to account for phylogenetic non-independence in analysis of among-species comparisons. Doughnut charts show the proportion of sample sizes (i.e., number of Pearson correlation coefficients) and the number of taxa.

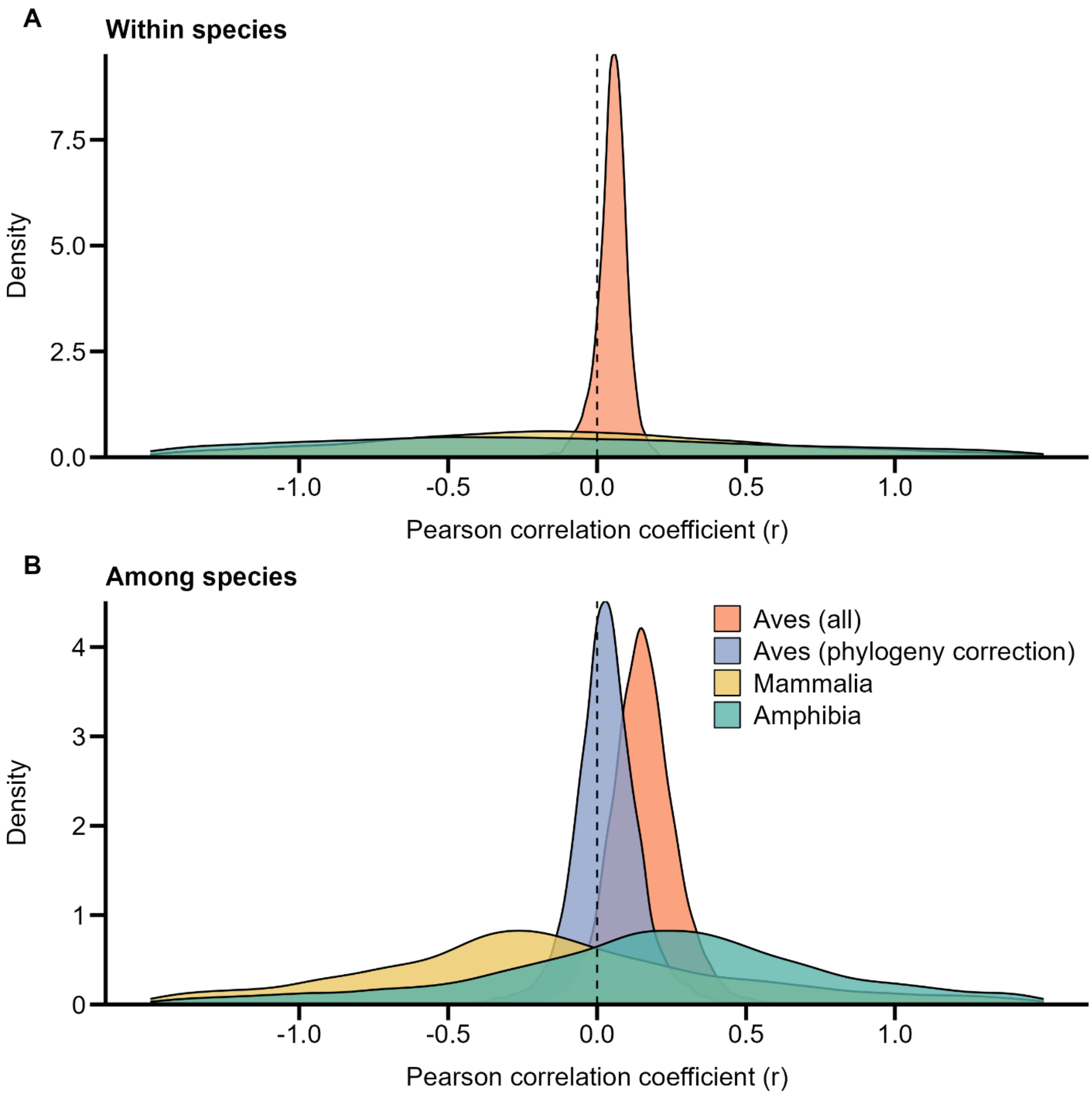


**Fig. S3.** Density plots showing posterior distributions obtained from Bayesian models for the differences in Pearson correlation coefficients (r) among animal classes obtained from general linear mixed-effects models (see Methods). Positive values indicate support for the AAH.

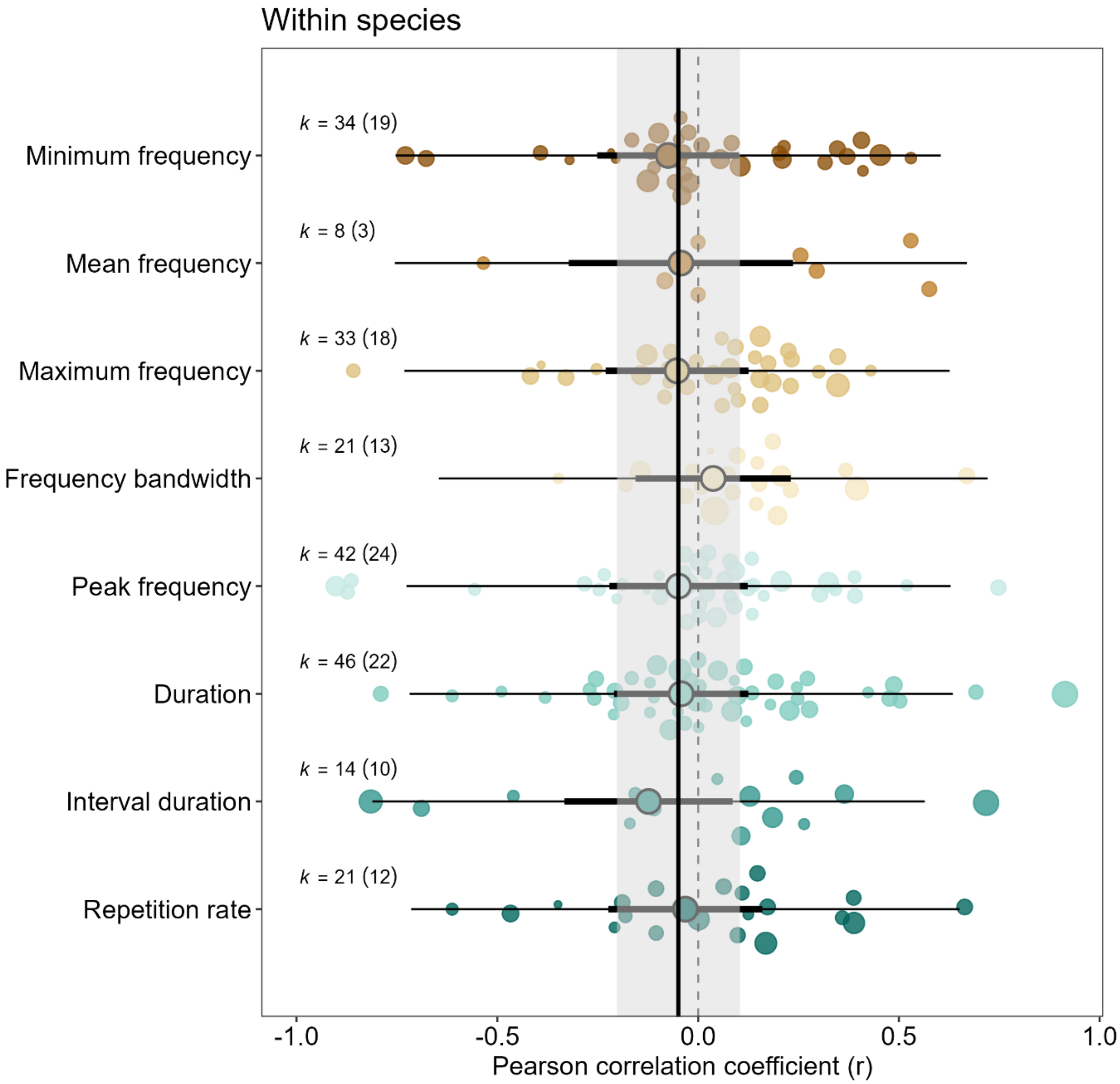


**Fig. S4.** Orchard plots for within-species comparisons showing individual effect sizes and mean effect size obtained from REML models for each acoustic variable. Number of effect sizes and number of taxa (k = number of effect sizes (number of taxa)) are shown on the left for each variable. Bold horizontal bars are 95% confidence intervals (CIs) and thin black lines are precision intervals. Positive Pearson correlation coefficients (r) indicate support for the acoustic adaptation hypothesis. Solid vertical line indicates the estimated global effect size obtained from the REML model with random factors only, and the grey area denotes the 95% CIs (*r* = -0.049, 95% CIs: -0.202 to 0.104). Circle size reflects effect size precision (log(1/SE)).

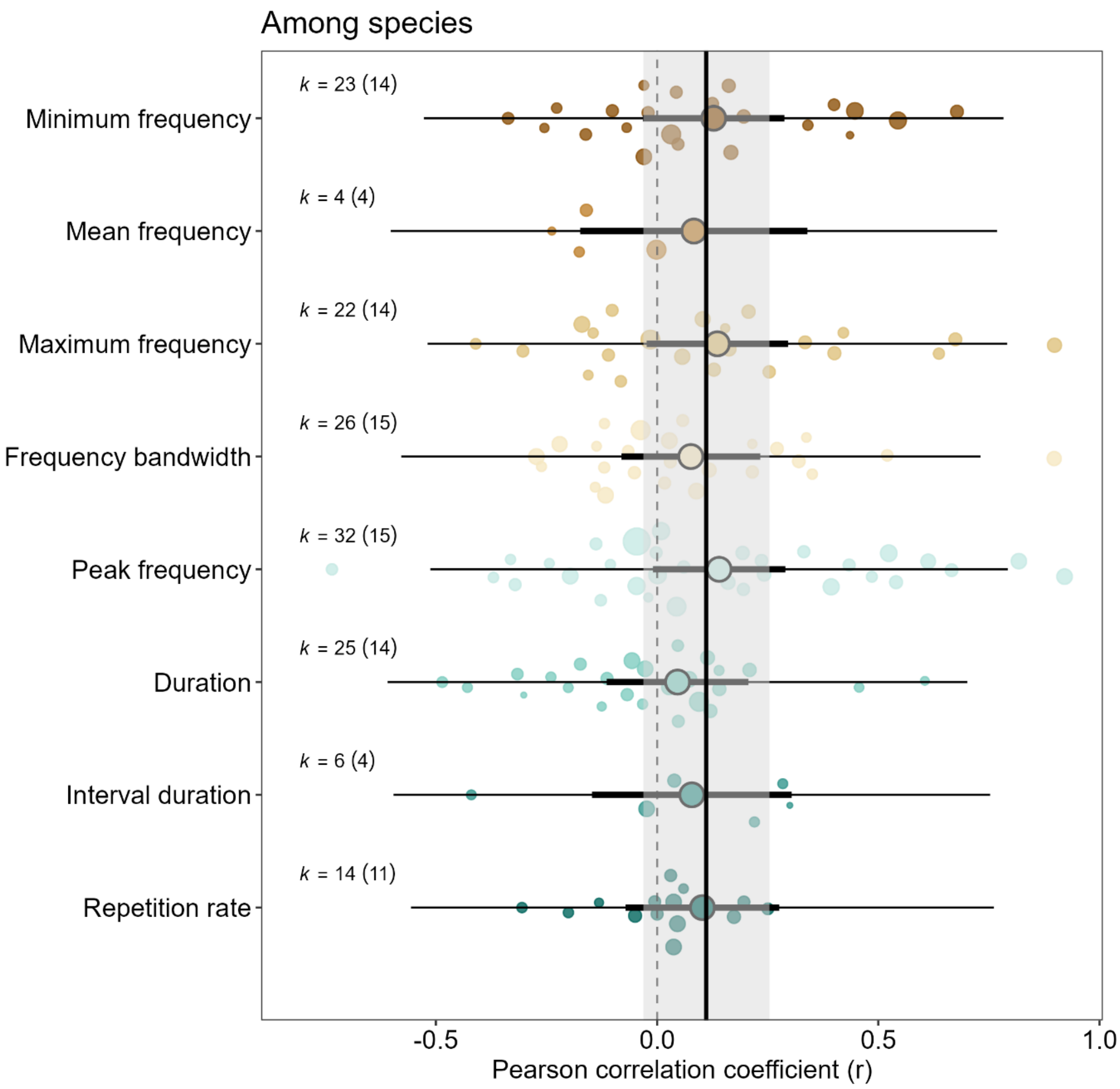


**Fig. S5.** Orchard plots for among-species comparisons showing individual effect sizes and mean effect size from REML models for each acoustic variable. Number of effect sizes and number of taxa (k = number of effect sizes (number of taxa)) are shown on the left for each variable. Bold horizontal bars are 95% confidence intervals (CIs) and thin black lines are precision intervals. Positive Pearson correlation coefficients (r) indicate support for the acoustic adaptation hypothesis. Solid vertical line indicates the estimated global effect size obtained from the REML model with random factors only, and the grey area denotes the 95% CIs (*r* = 0.111, 95% CIs: -0.031 to 0.254). Circle size reflects effect size precision (log(1/SE)).

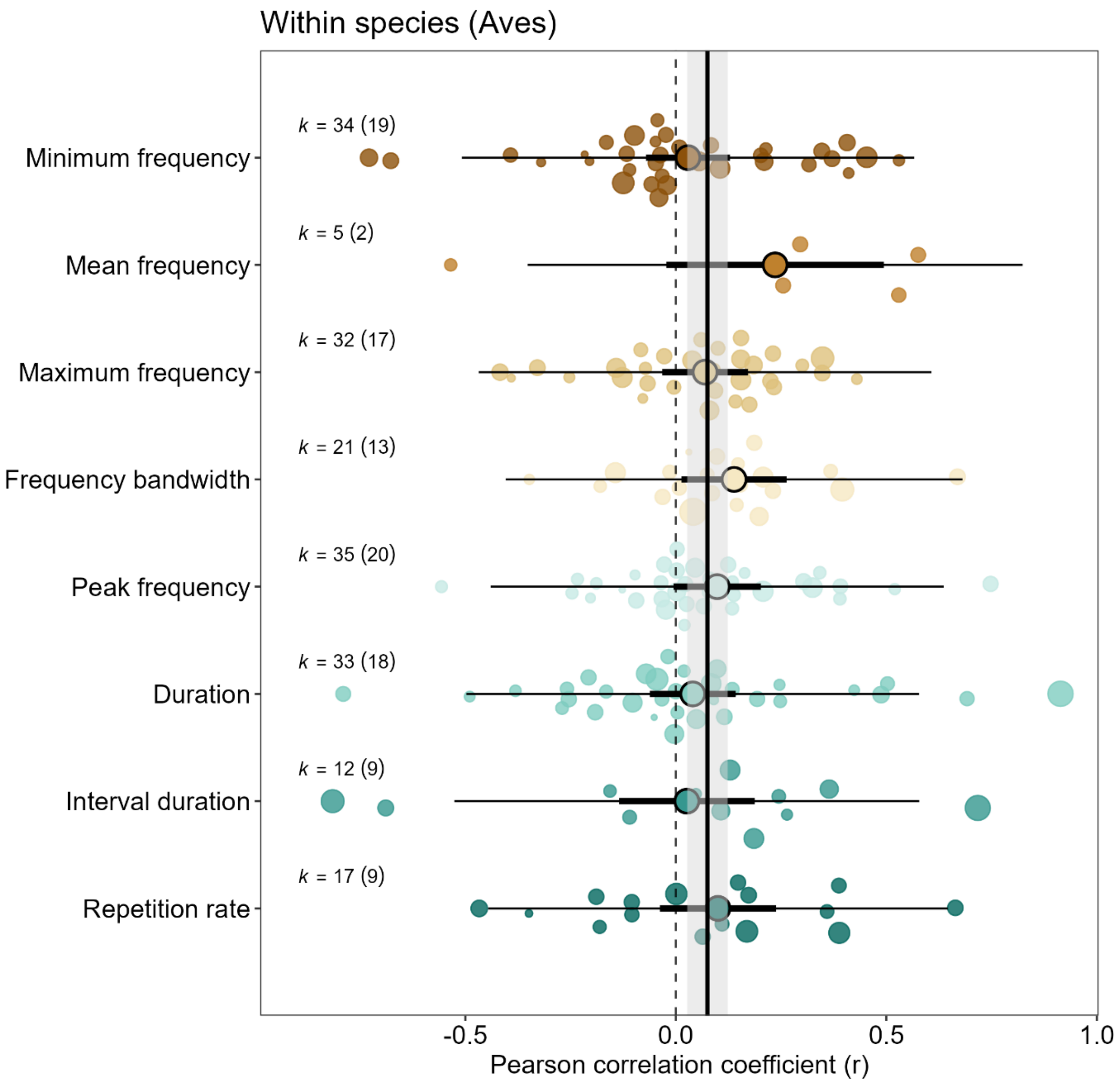


**Fig. S6.** Orchard plots for comparisons within bird species showing individual effect sizes and mean effect size obtained from REML models for each acoustic variable. Number of effect sizes and number of taxa (k = number of effect sizes (number of taxa)) are shown on the left for each variable. Bold horizontal bars are 95% confidence intervals (CIs) and thin black lines are precision intervals. Positive Pearson correlation coefficients (r) indicate support for the acoustic adaptation hypothesis. Solid vertical line indicates the estimated global effect size obtained from the REML model with random factors only, and the grey area denotes the 95% CIs (*r* = 0.075, 95% CIs: 0.027 to 0.123). Circle size reflects effect size precision (log(1/SE)).

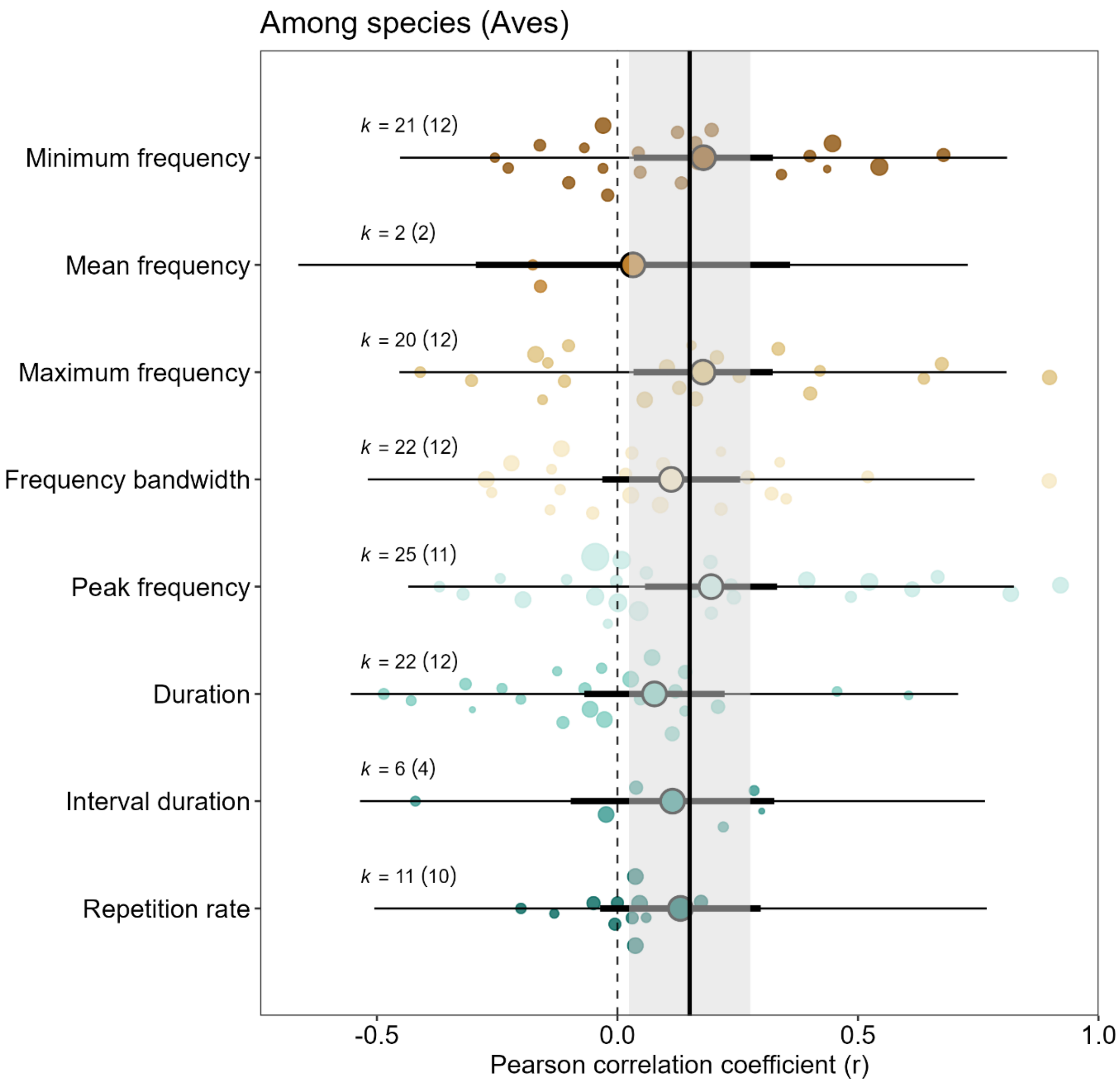


**Fig. S7.** Orchard plots for comparisons among bird species showing individual effect sizes and mean effect size from REML models for each acoustic variable. Number of effect sizes and number of taxa (k = number of effect sizes (number of taxa)) are shown on the left for each variable. Bold horizontal bars are 95% confidence intervals (CIs) and thin black lines are precision intervals. Positive Pearson correlation coefficients (r) indicate support for the acoustic adaptation hypothesis. Solid vertical line indicates the estimated global effect size obtained from the REML model with random factors only and the grey area denotes the 95% CIs (*r* = 0.150, 95% CIs: 0.024 to 0.276). Circle size reflects effect size precision (log(1/SE)).

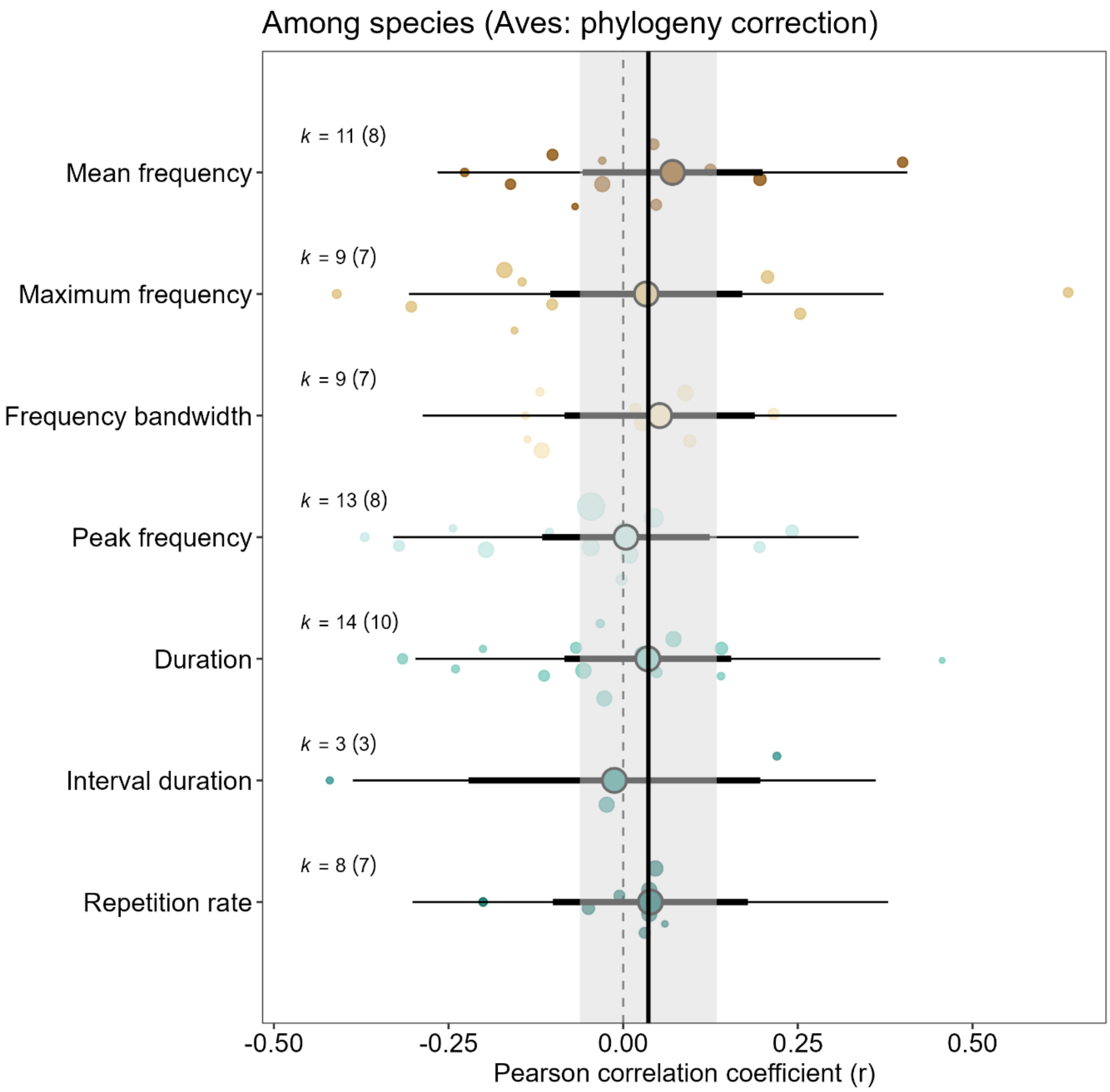


**Fig. S8.** Orchard plots for comparisons that accounted for phylogenetic non-independence (phylogeny correction) among bird species showing individual effect sizes and mean effect size from REML models for each acoustic variable. Number of effect sizes and number of taxa (k = number of effect sizes (number of taxa)) are shown on the left for each variable. Bold horizontal bars are 95% confidence intervals (CIs) and thin black lines are precision intervals. Positive Pearson correlation coefficients (r) indicate support for the acoustic adaptation hypothesis. Solid vertical line indicates the estimated global effect size from the REML model including only phylogeny corrected studies with random factors only, and the grey area denotes the 95% CIs (*r* = 0.036, 95% CIs: -0.062 to 0.134). Circle size reflects effect size precision (log(1/SE)).

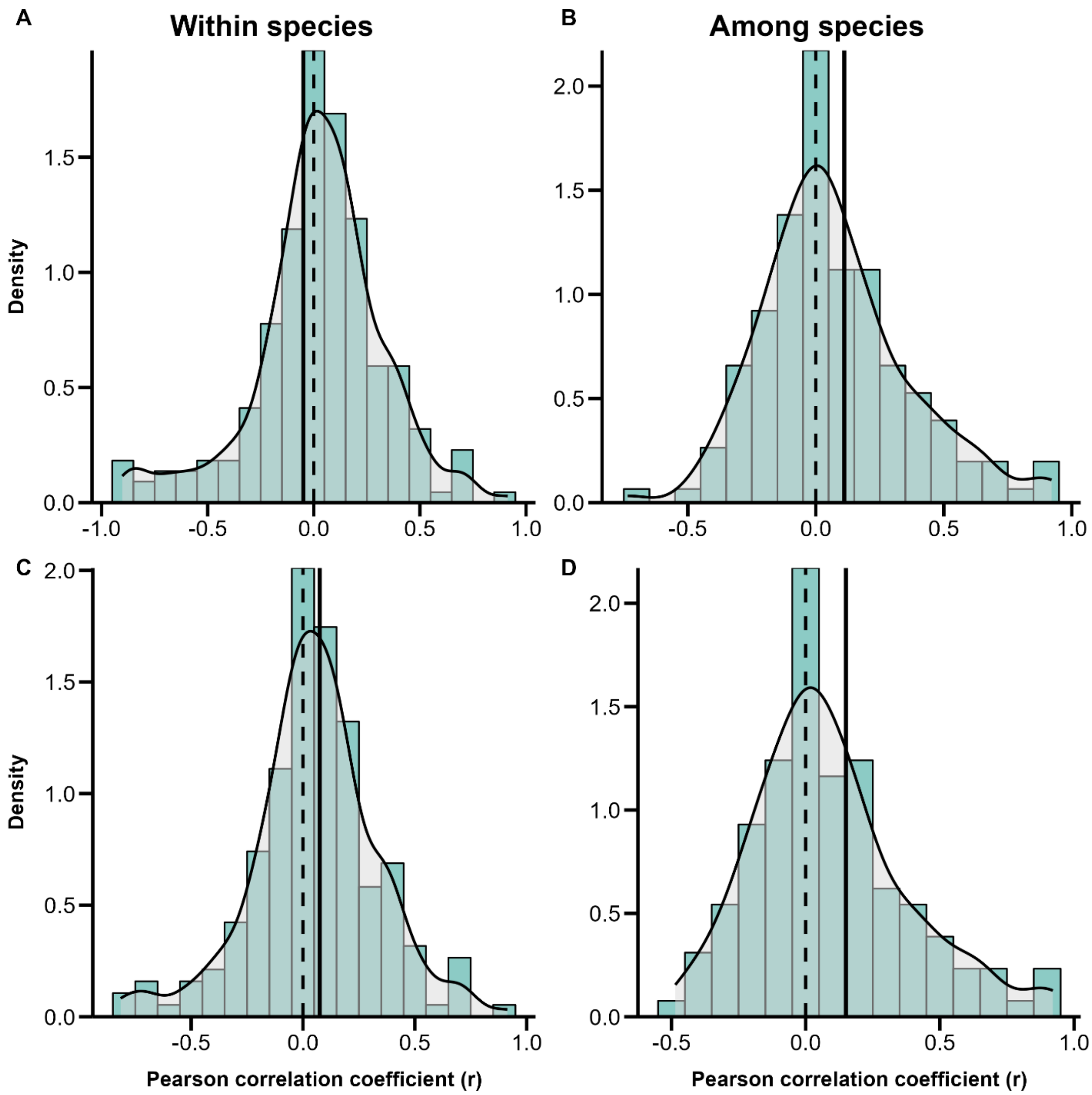


**Fig. S9.** Bar and density plots for within- (left) and among-species (right) comparisons showing the distribution of effect sizes (Pearson correlation coefficient (r)). Plots are shown for entire dataset (A and B) and for a subset of studies focusing on birds (C and D). Positive Pearson correlation coefficients (r) indicate support for the acoustic adaptation hypothesis. Solid vertical black lines indicate the estimated global effect size obtained from the REML model with random factors only for global models (A: r = -0.049; B: r = 0.111) and for the ones including only comparisons of birds (C: r = 0.075; D: r = 0.150).

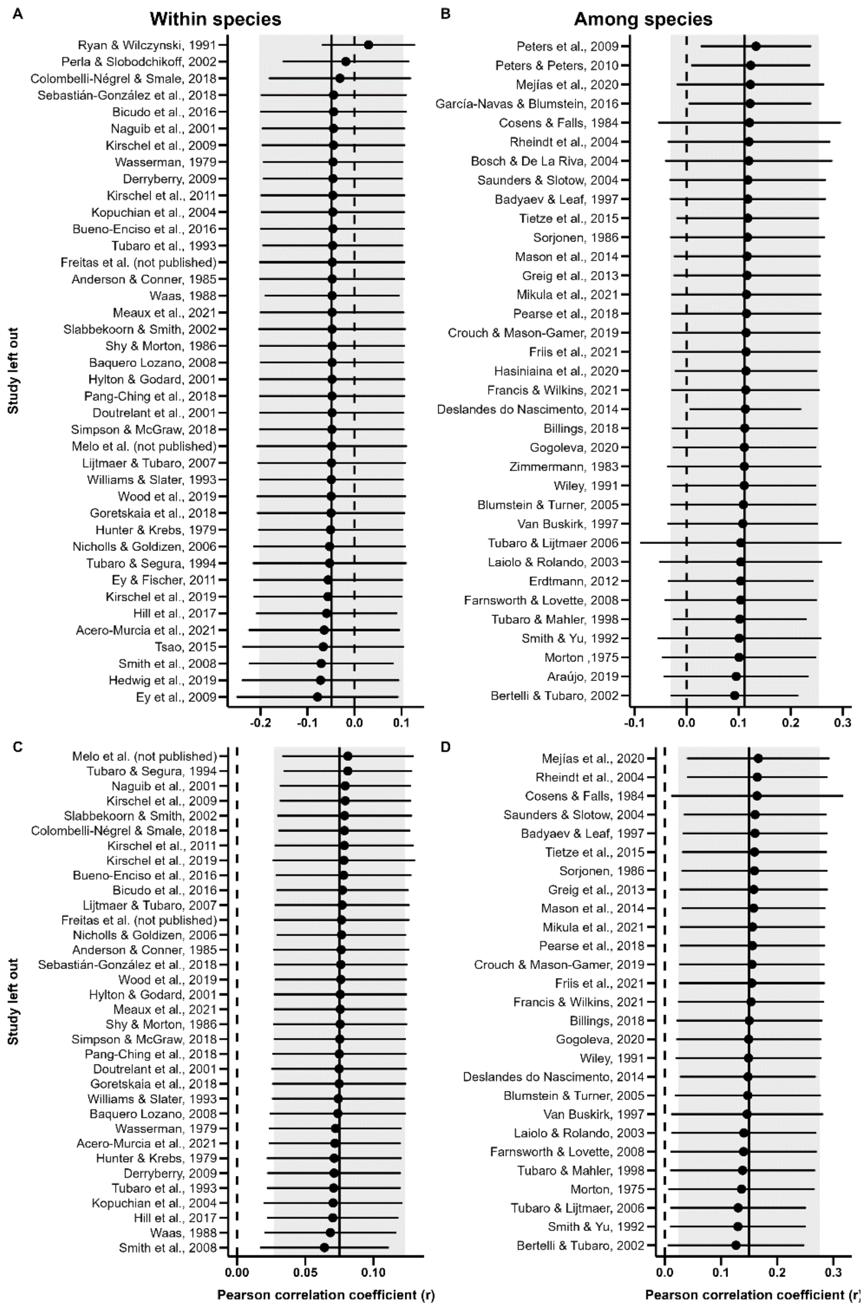


**Fig. S10.** Caterpillar plots of leave-one-out analyses for within- (left) and among-species (right) comparisons based on datasets with one study left out at a time from model fitting. Plots are shown for entire dataset (A and B) and for a subset of studies focusing on birds (C and D). Estimated global effect sizes are ordered by mean estimate, and with horizontal bars showing 95% confidence intervals (CIs). Positive Pearson correlation coefficients (r) indicate support for the acoustic adaptation hypothesis. Solid vertical black lines indicate the estimated global effect size obtained from the REML model with random factors only, and the grey area denotes the 95% CIs for global models (A: r = -0.049, 95% CIs: -

0.202 to 0.104; B: r = 0.111, 95% CIs: -0.031 to 0.254) and for the ones including only comparisons of birds (C: r = 0.075, 95% CIs: 0.027 to 0.123; D: r = 0.150, 95% CIs: 0.024 to 0.276).

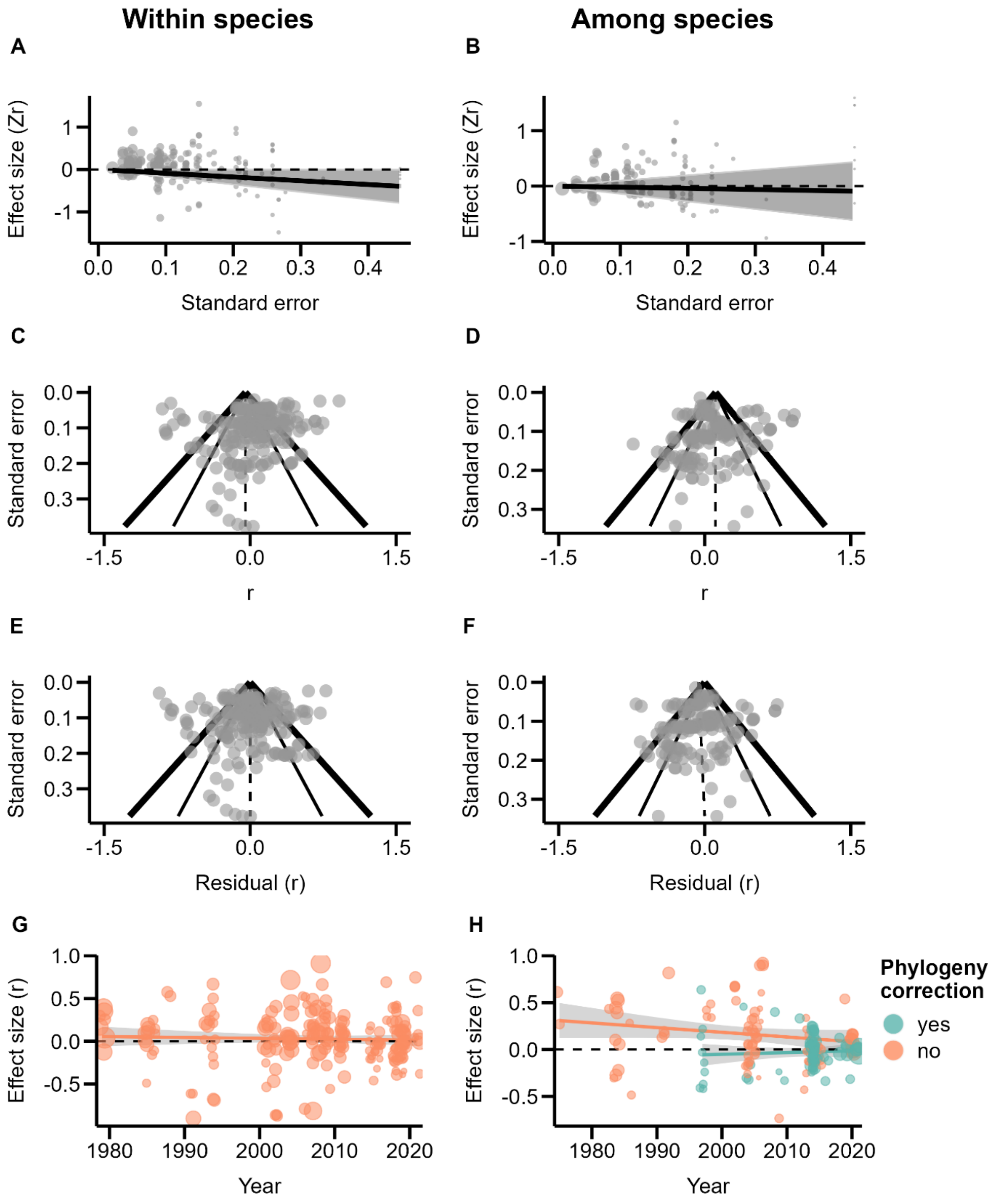


**Fig. S11**. Plots for the assessment of publication bias for within- (left) and among-species (right) comparisons. Regressions (A and B) show the relationship between effect size (Fisher's z) and its standard error testing whether studies with smaller sample sizes and lower precision are more likely to be published when reporting larger effect sizes. Solid lines show the fit of the multivariate linear mixed-effects model with year and all the moderators as fixed effects and study identifier, effect size identifier, taxon name, and phylogeny as random terms (within-species comparisons: slope = -0.887, 95% CI = [-

1.766, -0.008], t-value = -1.990, P-value = 0.048; among-species comparisons: slope = -0.206, 95% CI = [-1.382, 0.970], t-value = -0.346, P-value = 0.730). Circle size reflects effect size precision (log(1/SE)). Funnel plots for raw values (C and D) and for meta-analytic residuals (E and F) were obtained from multivariate linear mixed-effects model with all the moderators as fixed effects and study identifier, effect size identifier, taxon name, and phylogeny as random terms. Dashed lines indicate the estimated global effect size. Bold and thin black solid lines denote the expected 95% and 99% confidence limits, respectively, purely due to sampling heterogeneity. Asymmetries along the global effect size may reflect publication biases. Time-lag bias analysis (G and H) showing the relationship (linear regression) between effect size (Pearson correlation coefficient, r) and the year of publication of studies that did (and did not) account for phylogenetic non-independence (within-species comparisons: slope ± SE = -0.001 ± 0.002, t-value = -0.545, P-value = 0.586; among-species comparisons: slope[yes] ± SE = 0.002 ± 0.004, t-value = 0.514, P-value = 0.609, slope[no] ± SE = -0.005 ± 0.003, t-value = -1.625, P-value = 0.108). Circle size reflects effect size precision (1/SE).

**TABLES**

**Table S1**. Definition of each acoustic variable and the general predictions of the acoustic adaptation hypothesis for a closed habitat in comparison to an open habitat.

| Acoustic variable | Definition | Prediction for closed habitat |
|---|---|---|
| Minimum frequency | Lowest frequency of the signal | Lower (-) |
| Mean frequency | Average frequency of the signal | Lower (-) |
| Maximum frequency | Highest frequency of the signal | Lower (-) |
| Frequency bandwidth | Difference between the maximum and minimum frequencies of the signal | Lower (-) |
| Peak frequency | The frequency that contains most energy (highest amplitude) in the signal; also called dominant frequency | Narrower (-) |
| Duration | Length from start to end of the signal | Longer (+) |
| Interval duration | Duration of the time between consecutive signals, this is, the silent period or gap between signals | Longer (+) |
| Repetition rate | Number of signals emitted per unit of time | Lower (-) |

**Table S2**. Source data table of effect sizes included in the meta-analysis.

(provided in a separate file: AAH_meta_Sup_mat_Table_S2.csv)

**Table S3**. Number of studies and effect sizes for each moderator category included in this meta-analysis for primary studies of within- and among-species comparisons.

| | | **Within species** | | **Among species** | |
|---|---|---|---|---|---|
| **Moderator** | | **k Effect sizes** | **N Studies** | **k Effect sizes** | **N Studies** |
| Class | Aves | 189 | 34 | 129 | 27 |
| | Mammalia | 16 | 4 | 11 | 4 |
| | Amphibia | 14 | 2 | 12 | 4 |
| Acoustic variable | Minimum frequency | 34 | 22 | 23 | 17 |
| | Mean frequency | 8 | 4 | 4 | 4 |
| | Maximum frequency | 33 | 22 | 22 | 16 |
| | Frequency bandwidth | 21 | 14 | 26 | 17 |
| | Peak frequency | 42 | 22 | 32 | 24 |
| | Duration | 46 | 25 | 25 | 14 |
| | Interval duration | 14 | 13 | 6 | 6 |
| | Repetition rate | 21 | 14 | 14 | 8 |
| Habitat classification | Categorical | 119 | 25 | 115 | 29 |
| | Continuous | 100 | 15 | 37 | 6 |
| Habitat contrast* | 1 | 68 | 15[#] | 95 | 24 |
| | 0 | 51 | 11[#] | 20 | 5 |
| Body size correction | Corrected | 8 | 2 | 41 | 13 |
| | Not Corrected | 211 | 38 | 111 | 22 |
| Phylogeny correction | Corrected | 0 | 0 | 73 | 19 |
| | Not Corrected | 219 | 40 | 79 | 18 |
| Total | | 219 | 40 | 152 | 35 |

*Habitat contrast analysis was only scored for studies that employed a categorical classification of habitat type.
[#] Note that one primary study compared both more dissimilar habitats (contrast value = 1) and more similar habitats (contrast value = 0).

**Table S4**. Studies and effect sizes included in a previous meta-analysis testing the acoustic adaptation hypothesis in birds by Boncoraglio & Saino (2007) and effect sizes included in our study converted to Fisher's z. Inconsistencies between this and our study were divided in six different categories: (1) effect size differs more than 0.10; (2) effect size is similar but differs in sign; (3) effect size could not be retrieved for our study; (4) study excluded from our meta-analysis based on our selection criteria; (5) different methodological approach: authors might have pooled the data and we did not; (6) different methodological approach: we used raw values.

| Author(s) | Year | *Zr* effect size (Boncoraglio & Saino 2007) | | | | | *Zr* effect size (this study) | | | | |
|---|---|---|---|---|---|---|---|---|---|---|---|
| | | Maximum frequency | Minimum frequency | Frequency range | Peak frequency | Interval duration | Maximum frequency | Minimum frequency | Frequency range | Peak frequency | Interval duration |
| Anderson & Conner | 1985 | 0.166 | -0.040 | 0.200 | 0.002 | | 0.155 | -0.040 | 0.201 | 0.002 | |
| Badyaev & Leaf | 1997 | 0.436 [2] | 0.030 [2] | 0.141 [2] | 0.388 [2] | -0.224 | -0.436 [2] | -0.030 [2] | -0.141 [2] | -0.388 [2] | -0.224 |
| Bertelli & Tubaro | 2002 | 0.839 | 0.846 | 0.593 | 0.823 | | 0.818 | 0.825 | 0.577 | 0.803 | |
| Blumstein & Turner | 2005 | 0.268 | 0.07 | 0.252 | 0.166 | -0.034 | 0.348 | 0.134 | 0.332 | 0.241 | -0.039 |
| Cosens & Falls | 1984 | 0.057 | 0.750 [1] | -0.28 | 0.695 [1] | | 0.057 | 0.481 [1] | -0.280 | 0.416 [1] | |
| Cosens & Falls | 1984 | 0.143 | 0.639 | -0.394 [1] | 0.626 | | 0.103 | 0.610 | -0.224 [1] | 0.581 | |
| Doutrelant *et al.* | 2001 | 0.156 | -0.099 | 0.211 | | 0.335 [1] | 0.156 | -0.099 | 0.210 | | -0.188 [1] |
| Galeotti *et al.* | 1996 | 0.26 [4] | 0.075 [4] | 0.114 [4] | 0.235 [4] | 0.017 [4] | [4] | [4] | [4] | [4] | [4] |
| Hunter & Krebs | 1979 | 0.364 | -0.126 | 0.418 | | | 0.364 | -0.126 | 0.418 | | |
| Hylton & Godard | 2001 | -0.261 | -0.048 | | 0.165 | 0.271 | -0.259 | -0.048 | | 0.165 | -0.271 |
| Kopuchian *et al.* | 2004 | 0.049 | 0.062 | | 0.063 | 0.753 [1] | 0.039 | 0.055 | | 0.046 | -0.902 [1] |
| Morton | 1975 | | | 0.281 | 0.713 | | | | 0.278 | 0.713 | |
| Nicholls & Goldizen | 2006 | 0.46 | 0.59 | | 0.576 | | 0.460 | 0.590 | | 0.576 | |
| Nottebohm | 1975 | -0.530 [4] | -0.252 [4] | | | | [4] | [4] | | | |
| Payne | 1978 | -0.098 [4] | | | | 0.042 [4] | [4] | | | | [4] |
| Saunders & Slotow | 2004 | | | 0.375 [1] | | | | | -0.268 [1] | | |

| **Author(s)** | **Year** | ***Zr* effect size (Boncoraglio & Saino 2007)** | | | | | ***Zr* effect size (this study)** | | | | |
|---|---|---|---|---|---|---|---|---|---|---|---|
| | | **Maximum frequency** | **Minimum frequency** | **Frequency range** | **Peak frequency** | **Interval duration** | **Maximum frequency** | **Minimum frequency** | **Frequency range** | **Peak frequency** | **Interval duration** |
| Shy | 1983 | 0.921 [4] | 0.441 [4] | 0.628 [4] | | -0.224 [4] | 0.907 [4] | 0.433 [4] | 0.618 [4] | | 0.220 [4] |
| Shy & Morton | 1986 | 0.093 | -0.113 | 0.152 | | -0.161 [2] | 0.091 | -0.110 | 0.149 | | 0.158 [2] |
| Slabbekoorn & Smith | 2002 | -0.039 [5] | 0.310 [5] | | -0.029 [5] | -0.224 [5] | [5] | [5] | | [5] | [5] |
| Smith & Yu | 1992 | | | 0.783 [3] | 1.335 [1] | | | | [3] | 1.150 [1] | |
| Tubaro & Lijtmaer | 2006 | 1.594 | 0.469 | 1.463 | 1.73 [1] | 0.356 [2] | 1.463 [1] | 0.467 | 1.460 | 1.596 [1] | -0.310 [2] |
| Tubaro & Mahler | 1998 | -0.836 [6] | | -0.492 [6] | -0.492 [6] | | 0.449 [6] | | 0.366 [6] | 0.530 [6] | |
| Tubaro & Segura | 1994 | -0.067 | 0.222 | -0.185 | | -0.86 [2] | -0.072 | 0.217 | -0.182 | | 0.846 [2] |
| Tubaro *et al.* | 1993 | 0.081 | 0.516 [1] | | | 0.646 [1] | 0.187 | 0.213 [1] | | | -0.382 [1] |
| Waas | 1987 | | | | 0.446 [5] | | | | | [5] | |
| Wasserman | 1979 | | | | 0.218 [5] | | | | | [5] | |
| Wiley | 1991 | 0.130 | 0.165 | | 0.197 | 0.251 [3] | 0.129 | 0.163 | | 0.196 | [3] |

**Table S5**. Summary statistics of moderator models of within- and among-species comparisons using restricted maximum likelihood approach. Moderators include habitat classification (categorical versus continuous), habitat contrast of categorical classifications (0 versus 1), and body size correction (no versus yes). For comparisons among species, moderators also include phylogeny correction (no versus yes). The first category mentioned in parentheses is used as the reference level for the model results. Phylogenetically controlled multilevel meta-analytic single predictor models are shown with Omnibus tests ($Q_M$, Wald-type chi-square test) and pseudo-$R^2$.

| Type of study | Moderator | Estimate | ± SE | QM | P-value | $R^2$ | | |
|---|---|---|---|---|---|---|---|---|
| Within species | Habitat classification | -0.108 | ±0.084 | 1.663 | 0.197 | 0.002 | (0.000, | 0.511) |
| | Habitat contrast | -0.022 | ±0.097 | 0.05 | 0.823 | 0.000 | (0.000, | 0.323) |
| | Body size correction | -0.169 | ±0.197 | 0.738 | 0.390 | 0.000 | (0.000, | 0.392) |
| | Habitat classification (Aves) | -0.058 | ±0.051 | 1.281 | 0.258 | 0.002 | (0.000, | 0.569) |
| | Habitat contrast (Aves) | 0.004 | ±0.068 | 0.004 | 0.949 | 0.000 | (0.000, | 0.268) |
| | Body size correction (Aves) | -0.164 | ±0.121 | 1.832 | 0.176 | 0.007 | (0.000, | 0.385) |
| Among | Habitat classification | -0.033 | ±0.145 | 0.051 | 0.822 | 0.000 | (0.000, | 0.376) |
| | Habitat contrast | -0.255 | ±0.152 | 2.799 | 0.094 | 0.000 | (0.000, | 0.416) |
| | Body size correction | -0.159 | ±0.109 | 2.140 | 0.143 | 0.036 | (0.000, | 0.357) |
| | Phylogeny correction | -0.128 | ±0.083 | 2.355 | 0.125 | 0.088 | (0.000, | 0.452) |
| | Habitat classification (Aves) | -0.061 | ±0.149 | 0.171 | 0.679 | 0.000 | (0.000, | 0.382) |
| | Habitat contrast (Aves) | -0.268 | ±0.188 | 2.028 | 0.154 | 0.020 | (0.000, | 0.222) |
| | Body size correction (Aves) | -0.181 | ±0.122 | 2.201 | 0.138 | 0.049 | (0.000, | 0.368) |
| | Phylogeny correction | -0.191 | ±0.090 | 4.488 | 0.034 | 0.165 | (0.000, | 0.410) |

**Table S6**. Global tests using the Bayesian approach. Results of intercept-only general linear-mixed effects models are shown for the entire dataset (global model) and subsets with respect to animal class (Aves, Mammalia or Amphibia). For among-species comparisons of birds, the results are also shown for a subset of studies that accounted for phylogenetic non-independence (Aves (phylogeny correction)). Table shows number of effect sizes (k), number of species or taxonomic groups (N), estimates of r with 95% confidence intervals in parentheses and heterogeneity $I^2$ (variance arising from phylogenetic affinities, taxon, study identity, and observation). Model estimates are shown as posterior modes with 95% Highest Posterior Density (HPD) intervals in brackets.

| **Type of study** | **Taxonomic range** | **k** | **N** | **r** | **$I^2$Phylogeny** | **$I^2$Species** | **$I^2$Study** | **$I^2$Observation** |
|---|---|---|---|---|---|---|---|---|
| Within species | Global model | 219 | 37 | -0.031(-0.256,0.113) | 0.202(0.000, 0.361) | 0.160(0.000, 0.335) | 0.243(0.031,0.417) | 0.396(0.264, 0.529) |
| | Aves (all) | 189 | 32 | 0.058 (-0.068,0.145) | 0.131(0.000, 0.304) | 0.089(0.000, 0.211) | 0.116(0.000,0.266) | 0.663(0.439, 0.882) |
| | Mammalia | 16 | 3 | -0.148(-2.228,2.189) | 0.368(0.000, 0.744) | 0.351(0.000, 0.725) | 0.189(0.000,0.541) | 0.091(0.014, 0.204) |
| | Amphibia | 14 | 2 | -0.404(-3.858,3.176) | 0.320(0.000, 0.707) | 0.321(0.000, 0.692) | 0.324(0.000,0.701) | 0.035(0.000, 0.099) |
| Among species | Global model | 152 | 20 | 0.102 (-0.134,0.377) | 0.231(0.000, 0.443) | 0.140(0.000, 0.292) | 0.401(0.242,0.581) | 0.229(0.144, 0.324) |
| | Aves (all) | 129 | 14 | 0.147 (-0.059,0.377) | 0.181(0.000, 0.388) | 0.142(0.000, 0.298) | 0.435(0.266,0.601) | 0.242(0.144, 0.344) |
| | Aves (phylogeny correction) | 67 | 11 | 0.027 (-0.189,0.259) | 0.254(0.000, 0.541) | 0.303(0.020, 0.545) | 0.260(0.039,0.486) | 0.182(0.039, 0.349) |
| | Mammalia | 11 | 3 | -0.260(-2.239,1.798) | 0.365(0.001, 0.744) | 0.347(0.000, 0.723) | 0.246(0.000,0.601) | 0.042(0.000, 0.131) |
| | Amphibia | 12 | 3 | 0.227 (-1.829,2.132) | 0.349(0.000, 0.709) | 0.327(0.001, 0.691) | 0.190(0.000,0.514) | 0.135(0.011, 0.341) |

**Table S7**. Test of moderators explaining for within- and among-species comparisons using the Bayesian approach. Moderators include habitat classification (categorical versus continuous), habitat contrast of categorical classifications (0 versus 1), and body size correction (no versus yes). For comparisons among species, moderators also include phylogeny correction (no versus yes). The first category mentioned in parentheses is used as the reference level for the model results. Model estimates (i.e., estimated difference between groups) are shown as posterior modes with 95% Highest Posterior Density (HPD) intervals obtained from phylogenetically controlled general linear-mixed effects models. The variance explained by the entire model and by the fixed factors is given as the conditional $R^2$ and the marginal $R^2$, respectively, with 95% HPD intervals in brackets.

| Type of study | Moderator | Estimate | $R^2$ (conditional) | $R^2$ (marginal) |
|---|---|---|---|---|
| Within species | Habitat classification | -0.003(-0.275, 0.052) | 0.856(0.821, 0.889) | 0.044(0.000, 0.132) |
| | Habitat contrast | -0.030(-0.257, 0.179) | 0.914(0.885, 0.940) | 0.028(0.000, 0.101) |
| | Body size correction | -0.044(-0.565, 0.202) | 0.855(0.819, 0.888) | 0.022(0.000, 0.078) |
| | Habitat classification (Aves) | 0.084(-0.180, 0.046) | 0.838(0.795, 0.880) | 0.023(0.000, 0.076) |
| | Habitat contrast (Aves) | 0.072(-0.251, 0.257) | 0.911(0.879, 0.943) | 0.033(0.000, 0.126) |
| | Body size correction (Aves) | 0.061(-0.425, 0.090) | 0.837(0.795, 0.879) | 0.021(0.000, 0.068) |
| Among species | Habitat classification | 0.119(-0.341, 0.236) | 0.787(0.741, 0.835) | 0.046(0.000, 0.164) |
| | Habitat contrast | 0.316(-0.570, 0.068) | 0.813(0.763, 0.863) | 0.100(0.000, 0.246) |
| | Body size correction | 0.151(-0.353, 0.069) | 0.785(0.738, 0.833) | 0.063(0.000, 0.174) |
| | Phylogeny correction | 0.168(-0.307, 0.034) | 0.786(0.738, 0.835) | 0.078(0.000, 0.208) |
| | Habitat classification (Aves) | 0.174(-0.375, 0.219) | 0.782(0.731, 0.832) | 0.059(0.000, 0.203) |
| | Habitat contrast (Aves) | 0.398(-0.667, 0.095) | 0.809(0.754, 0.867) | 0.114(0.000, 0.284) |
| | Body size correction (Aves) | 0.206(-0.412, 0.087) | 0.780(0.730, 0.834) | 0.084(0.000, 0.219) |
| | Phylogeny correction (Aves) | 0.252(-0.382, -0.012) | 0.782(0.730, 0.834) | 0.129(0.000, 0.287) |

**Table S8**. Sensitivity analysis examining the robustness of the overall effect size (r) to the assumption of different ρ values (within-study correlation) used to construct the variance-covariance metrics that account for non-independence of sampling variances. Table shows estimates of r with 95% confidence intervals in parentheses.

| Type of study | Taxonomic range | ρ | r | | | z-value | P-value |
|---|---|---|---|---|---|---|---|
| Within species | Global model | 0.3 | -0.048 | (-0.198, | 0.103) | -0.622 | 0.534 |
| | | 0.5 | -0.049 | (-0.202, | 0.104) | -0.622 | 0.534 |
| | | 0.7 | -0.049 | (-0.205, | 0.106) | -0.624 | 0.533 |
| | | 0.9 | -0.05 | (-0.208, | 0.107) | -0.625 | 0.532 |
| | Aves (all) | 0.3 | 0.072 | (0.026, | 0.118) | 3.073 | 0.002 |
| | | 0.5 | 0.075 | (0.027, | 0.123) | 3.057 | 0.002 |
| | | 0.7 | 0.078 | (0.027, | 0.128) | 3.029 | 0.002 |
| | | 0.9 | 0.079 | (0.028, | 0.131) | 2.998 | 0.003 |
| Among species | Global model | 0.3 | 0.109 | (-0.048, | 0.266) | 1.358 | 0.175 |
| | | 0.5 | 0.111 | (-0.031, | 0.254) | 1.532 | 0.126 |
| | | 0.7 | 0.115 | (-0.011, | 0.241) | 1.794 | 0.073 |
| | | 0.9 | 0.12 | (0.012, | 0.227) | 2.186 | 0.029 |
| | Aves (all) | 0.3 | 0.146 | (0.017, | 0.274) | 2.221 | 0.026 |
| | | 0.5 | 0.15 | (0.024, | 0.276) | 2.331 | 0.02 |
| | | 0.7 | 0.153 | (0.031, | 0.276) | 2.454 | 0.014 |
| | | 0.9 | 0.156 | (0.035, | 0.276) | 2.537 | 0.011 |
| | Aves (phylogeny correction) | 0.3 | 0.036 | (-0.067, | 0.139) | 0.679 | 0.497 |
| | | 0.5 | 0.036 | (-0.062, | 0.134) | 0.724 | 0.469 |
| | | 0.7 | 0.038 | (-0.064, | 0.140) | 0.733 | 0.463 |
| | | 0.9 | 0.038 | (-0.067, | 0.143) | 0.714 | 0.475 |

**Table S9.** Global effect sizes (Fisher's z) for the acoustic adaptation hypothesis obtained from the restricted maximum likelihood approach. Results of intercept-only General Linear-Mixed Effects Models are shown for the entire dataset (global model) and for subsets with respect to animal class (Aves). For among-species comparisons of birds, the results are shown for a subset of studies that accounted for phylogenetic non-independence (Aves (phylogeny correction)). Table shows number of effect sizes (k), number of species or taxa (N), and estimates of Fisher's z and the 95% confidence intervals in parentheses.

| Type of study | Dataset | k | N | z | | | z-value | P-value |
|---|---|---|---|---|---|---|---|---|
| Within species | Global model | 219 | 37 | 0.036 | (-0.049, | 0.120) | 0.829 | 0.407 |
| | Aves (all) | 189 | 32 | 0.081 | (0.028, | 0.134) | 3.004 | 0.003 |
| Among species | Global model | 152 | 20 | 0.115 | (-0.048, | 0.278) | 1.388 | 0.165 |
| | Aves (all) | 129 | 14 | 0.136 | (0.023, | 0.249) | 2.358 | 0.018 |
| | Aves (phylogeny correction) | 67 | 11 | 0.028 | (-0.068, | 0.124) | 0.572 | 0.568 |
